# Scalable and rare-variant aware genome inference across the 1kGP cohort

**DOI:** 10.64898/2026.06.29.735275

**Authors:** Jana Ebler, Timofey Prodanov, Andrew Blair, Samuel K. Lee, Peter Ebert, Human Pangenome Reference Consortium, Benedict Paten, Tobias Marschall

**Author notes:** Contributing authors.

## Abstract

Pangenome graphs built from haplotype-resolved *de novo* assemblies enable accurate analysis of genetic variation. The short-read-based tool PanGenie efficiently genotypes variants discovered in a pangenome across large cohorts and outperforms linear reference-based methods for structural variants (SVs). However, it cannot detect novel variants absent from the graph, missing many rare SVs (allele frequency ***<* 1%**) and was limited to graphs with 254 haplotypes. First, we introduce a haplotype sampling step that reduces the number of haplotypes using sample-specific k-mers before genotyping, decreasing runtime twelvefold and memory usage 1.4-fold at 30x coverage. Second, we present a polishing workflow that corrects residual errors in haplotypes inferred from PanGenie genotypes and incorporates rare and private mutations. We genotype 3,202 samples from the 1000 Genomes Project and use low-coverage ONT data (967 samples) for polishing. We achieve a median QV of 46 and provide the 1,934 polished haplotype sequences as a community resource.

## 1 Introduction

Advances in long-read sequencing technologies [1] as well as novel methods for haplotype-resolved assembly [2–5], enable near telomere-to-telomere *de novo* assembly of haplotypes [3, 6, 7]. Such haplotypes enable the construction of pangenome graphs [6, 8], which combine multiple reference genomes and, unlike a single linear reference, can capture genetic variability of a species. The Human Pangenome Reference consortium has recently generated a large pangenome for humans, including 462 near telomere-to-telomere assembled haplotypes.

Pangenomes have been demonstrated to improve downstream analyses [6, 8, 9], including genotyping which describes the process of determining whether both, one or none of the chromosomal copies of a diploid individual carry a specific variant allele [9]. Short reads remain widely used due to their low cost and the availability of large public databases [10–15]. Traditional short-read based approaches relying on a linear reference genome miss more than half of all SVs carried by a human individual [16], making it impossible to study these missed variants in downstream analyses. This is critical since SVs affect more bases than small variants and are thus a major contributor to phenotypic variation and diseases [10, 17, 18].

More recent genotyping approaches use pangenomes instead of a linear reference. VG’s short-read mapper Giraffe genotypes SVs based on alignments to a pangenome graph, reducing reference bias [19]. However, read alignment is computationally expensive. We have introduced PanGenie [9], a short-read genotyper using a pangenome in combination with counts of allele specific k-mers derived from the sequencing reads. Similarly, KAGE [20, 21] is a k-mer based method designed for pangenomes with larger numbers of haplotypes. Since PanGenie and KAGE do not rely on read alignment, they are much faster compared to alignment-based tools [9, 20].

Genotypers typically output unphased genotypes but can be combined with population-based phasing tools like SHAPEIT [22] or GLIMPSE [23, 24] to construct consensus haplotypes by implanting phased variant alleles into the underlying reference genome. However, genotypers only consider known variants and cannot detect novel variant alleles not represented in the underlying pangenome reference. Therefore, corresponding regions in the consensus haplotypes are inaccurate. Several tools exist for polishing haplotype sequences based on short reads [25–27] or long reads [28– 31] aiming to detect inconsistencies between reads and haplotype sequences. However, these methods mostly focus on single-base or small indel errors, and/or assume reads to be separated by haplotype already, which is not a trivial task for diploid genomes.

In this paper, we propose a new version of PanGenie, PanGenie v4, that includes a preprocessing step constructing a small, sample-specific set of haplotypes from the full pangenome graph. This allows more efficient genotyping on graphs with several hundreds of haplotypes and increases scalability up to biobank scale. Furthermore, we propose a polishing workflow to refine haplotypes derived from PanGenie genotypes based on short and low coverage long reads, correcting errors and including (rare) variants missed from the pangenome. We apply our methods to genotype 3,202 samples of the 1000 Genomes Project (1kGP) and generate polished haplotype sequences for a subset of 967 samples for which additional low coverage ONT data is available.

## 2 Results

### 2.1 Pangenome graph

The Human Pangenome Reference Consortium (HPRC) created their Release 2 pangenome graph from 462 near telomere-to-telomere assembled human haplotypes (referred to HPRC2 in this manuscript). The graph was constructed with Minigraph-Cactus [8] and uses the CHM13 reference genome as a backbone. Graph bubbles were called from the graph and decomposed into nested alleles using an approach we had introduced previously [6] (Methods). Using this approach, we have identified 36,483,743 SNP alleles, 10,512,438 indel alleles (*<* 50 bp), 127,964 SV deletion alleles, 643,535 SV insertion alleles and 129,530 other SV alleles. Here, deletions include all variants for which the length of the reference allele is *≥* 50 and the length of the alternative allele is 1. Insertions are defined vice versa. All other variants for which either the reference or alternative allele is *≥* 50 bp are defined as “other” SVs. Variant calls are relative to the CHM13 reference genome.

### 2.2 Haplotype sampling

We have previously introduced our tool PanGenie [9], which uses a pangenome reference in combination with short-read sequencing data to genotype genetic variants. PanGenie determines unique k-mers for each variant bubble in the graph and counts them in the sequencing reads of the sample to be genotyped (Figure 1a). It uses these counts in combination with haplotype path information inherent in the pangenome graph to derive a genotype for each bubble [9].

**Fig. 1.**
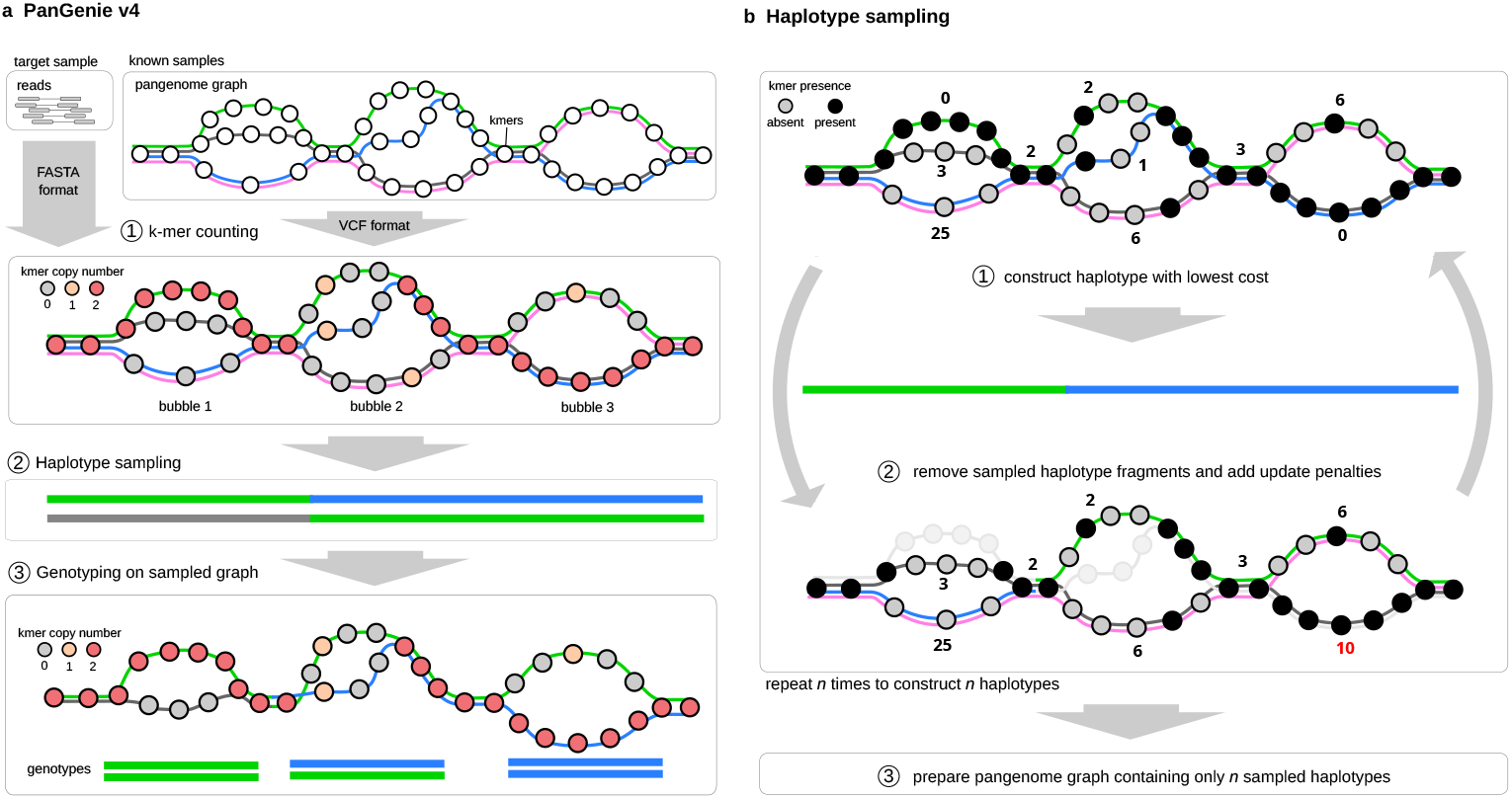
Haplotype sampling. **a** The input to PanGenie v4 are short read sequencing reads of the sample to be genotyped, as well as a VCF file encoding bubbles of a pangenome graph. PanGenie defines a set of unique k-mers for each bubble and counts them in the sequencing data (Step 1). It then constructs a smaller set of haplotype paths using the new haplotype sampling algorithm (Step 2) and proceeds with genotyping based on only these haplotypes instead of the full panel (Step 3). **b** In each step of the haplotype sampling algorithm, a new haplotype path is constructed based on coverages of unique k-mers in the reads. Already sampled path segments are removed from the graph and penalties are updated for already covered alleles prior to the next iteration.

Because the original method did not scale well with increasing numbers of haplotypes, previous versions of PanGenie accelerated genotyping by partitioning the set of haplotype paths into smaller subsets. Genotyping was performed independently on each chunk, producing genotype likelihoods for all bubbles based on the corresponding haplotype subset, which were subsequently combined to obtain final genotype predictions [3]. However, this approach does not scale well to the size of the HPRC2 graph. To address these limitations, we introduce a new haplotype sampling step which constructs a smaller set of paths from the full set of haplotypes represented in the graph. Each such haplotype path is a mosaic of the original set of haplotypes and is constructed in a sample specific manner by selecting path segments that share unique k-mers with the target sample. The objective is to minimize the number of recombinations needed to construct each path from the full set of input haplotypes, while maximizing the number of covered unique k-mers present in the sequencing reads.

We have developed a heuristic algorithm which creates new haplotype paths based on a dynamic programming (DP) approach (Figure 1b). In each iteration, the algorithm creates a DP table, consisting of one row per haplotype in the pangenome and one column per variant bubble. For each unique k-mer of each bubble, we determine whether it is present or absent in the sequencing reads of the individual to be genotyped. We then assign a local cost to each haplotype based on the fraction of unique k-mers it shares with the sample (Methods). The more unique k-mers are shared, the lower the cost. We furthermore define switch costs between columns based on recombination probabilities derived from the Li-Stephens model [32] (Methods). Then, we construct a haplotype path with the minimal overall costs through backtracing. Before generating the next haplotype using the same approach, we add penalties to all DP table cells traversed by the minimal path to encourage the algorithm to select different path segments in the next iteration. By repeating this process for *i* iterations, we construct *i* new haplotype mosaics. We then proceed by genotyping based on only the constructed haplotype paths instead of the full set of paths in the graph using the PanGenie algorithm.

### 2.3 Benchmarking of PanGenie v4

#### 2.3.1 Leave-one-out experiments

We tested the genotyping performance of PanGenie v4 with different parameters for the number of haplotypes sampled in the newly implemented haplotype sampling step. For this purpose, we performed “leave-one-out” experiments in which we repeatedly remove one sample from the panel VCF derived from the HPRC2 graph and use the remaining haplotypes as input to PanGenie to genotype the left-out sample from short-read data of the same sample. The assembly based genotypes of the left-out sample are used as a ground truth for evaluation. We use the weighted genotype concordance (wGC) [9] and F-score statistics computed with RTG vcfeval [33] and Truvari [34] as evaluation metrics (Supplementary Material). We ran PanGenie v4 with sampling sizes varying between 5 and 25 (Figure 2a, Supplementary Figure 2 and 3). For comparison, we also ran it without sampling using the chunking strategy implemented by previous versions, which independently ran genotyping on subsets of 108 haplotype paths and combined resulting genotype likelihoods (“PanGenie v4 chunking”). In general, PanGenie v4 performs better in most cases than PanGenie v4 with chunking, and genotyping performances tend to increase across all variant types as the number of sampled haplotypes is increased (Figure 2a, Supplementary Figures 2 and 3). At the same time, PanGenie v4 with the sampling step is around 12*×* faster (single-core) compared to PanGenie v4 with chunking and uses less memory (Figure 2b and c, Supplementary Figure 4). With 30*×* Illumina data used for our experiments, PanGenie v4 runs in around 55 mins using around 53 GB of RAM with 24 cores (Figure 2b and c) with 15 sampled haplotypes. Based on these results, we set the default number of sampled haplotypes to 15, as it provides high genotyping accuracy and reasonably low RAM usage. All results shown in the following use this default value.

**Fig. 2.**
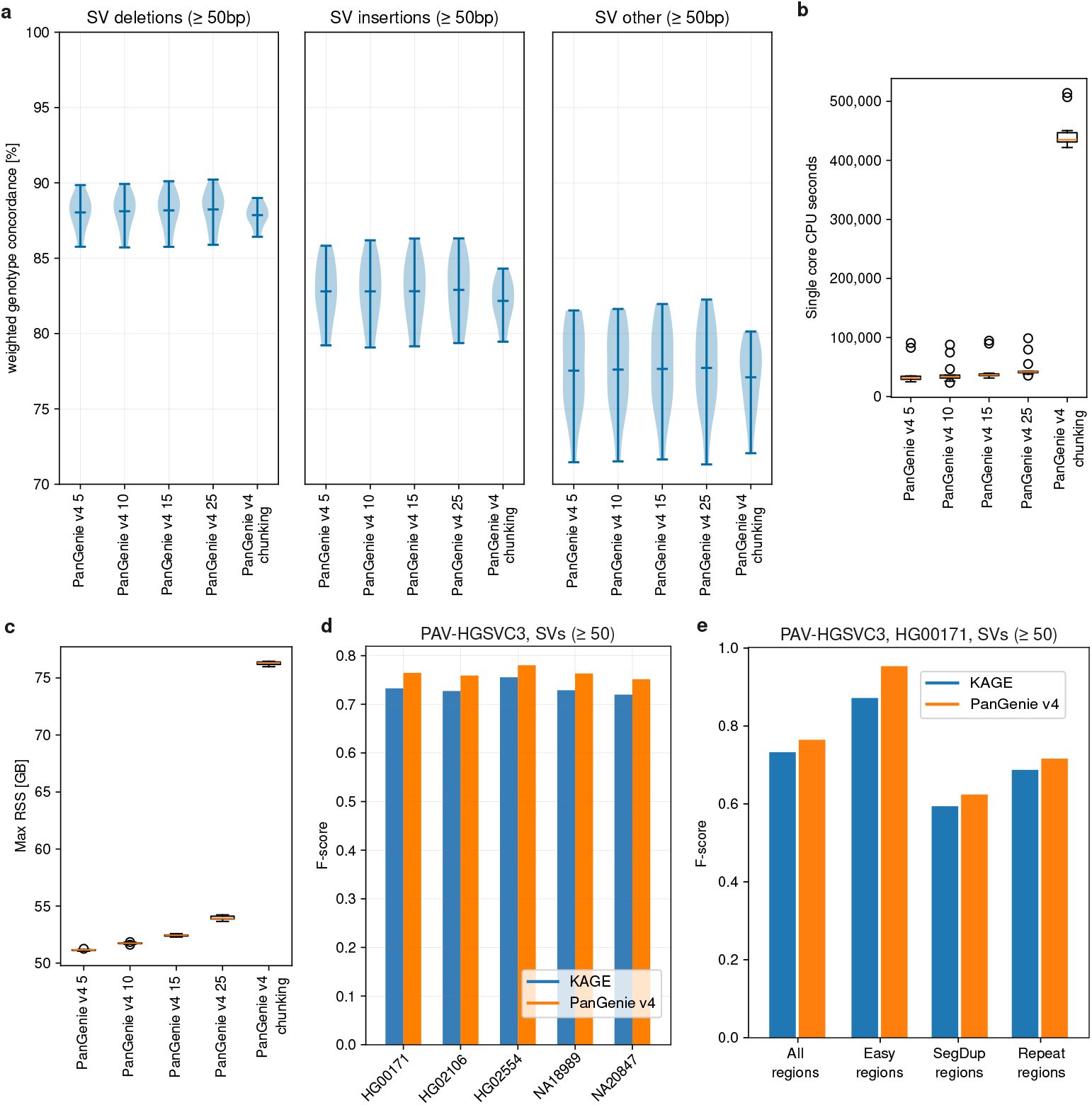
Benchmarking of genotyping tools. **a** Leave-one-out experiment with PanGenie v4 with different numbers of sampled haplotypes and with chunking on HPRC2 data, evaluated across 13 samples. For each sample, we removed its haplotypes from the input pangenome prior to genotyping and used it as ground truth for evaluation. **b** CPU seconds (single-core) of PanGenie v4 with different numbers of sampled haplotypes and PanGenie v4 with chunking. **c** Peak memory usage (GB) of PanGenie v4 with different sampling sizes and PanGenie v4 with chunking, run with 24 cores. **d** F-scores computed with truvari for PanGenie v4 and KAGE genotypes based on the assembly-based HGSVC3 PAV ground truth SVs for 5 evaluation samples. **e** F-scores computed with truvari for PanGenie v4 and KAGE genotypes based on the assembly-based HGSVC3 PAV ground truth SVs for HG00171 stratified by genomic regions defined by GIAB.

We furthermore repeated the same leave-one-out experiment using the previously published HGSVC3 pangenome graph instead of the HPRC2 graph. The HGSVC3 graph contained 216 human haplotypes and was constructed using the same Minigraph-Cactus pipeline [7]. Overall, we observed higher genotying accuracies when using the HPRC2 graph (Supplementary Figure 5). This can be explained by the larger number of haplotypes represented in the HPRC2 graph and the higher quality of the assemblies used to construct this graph.

#### 2.3.2 Comparison to other genotyping methods

We have previously demonstrated that PanGenie outperforms linear reference-based methods in terms of speed and genotyping accuracy, especially for SVs [3, 9]. In the meantime, another pangenome-based short-read genotyper, KAGE [20, 21], was introduced and recently used by the *All Of Us* consortium to genotype a large cohort of human samples [35]. We therefore compared our genotyping method to KAGE.

Due to issues running the latest version of KAGE2 (Supplementary Material), we ran a slightly modified version of KAGE v0.1.1 previously developed by the *All of Us* consortium [35], which included minor changes to improve scalability and robustness (Supplementary Material). This version relies on GLIMPSE1[23] for phasing resulting genotypes. All KAGE results described below refer to this modified version of KAGE. We genotyped five independent samples (HG00171, NA18989, NA20847, HG02106, HG02554), which are not in the HPRC2 graph, using PanGenie v4 and KAGE and evaluated the resulting genotypes based on independent ground truth SV calls produced by the HGSVC [7]. These ground truth sets included a PAV callset generated directly from assemblies of these samples as well as a Minigraph-Cactus callset generated from a pangenome graph of the HGSVC assemblies [7]. We used Truvari [34] in order to evaluate genotyping results. Overall, PanGenie performed slightly better than KAGE for all samples and regions evaluated (Figure 2 d and e). Across all autosomes, PanGenie produced F-scores between 0.73 and 0.78 for SVs. These numbers were between 3.2–4.7 % higher than for KAGE (Figure 2 d, Supplementary Figure 8). For SNPs and indels, PanGenie improved by 1.8–2.8% over KAGE across all five samples (Supplementary Figure 6). We furthermore stratified genotyping results by different regions defined by Genome in a Bottle (GIAB, v3.3), including “Easy” regions that are well accessible by short reads, “SegDup” including regions with segmental duplications and “Repeat” regions, including tandem repeats and homopolymers. PanGenie performed better across all regions tested, especially in “Easy” regions, in which it reached an F-score of 0.95 for sample HG00171 when restricting to SVs (improvement of 9.4% over KAGE). Numbers were similar for the other four samples evaluated (Supplementary Figure 8). For SNPs and indels, improvements were largest for segmental duplications (8.8–12.4% improvements for the five samples evaluated, Supplementary Figure 6). When using the Minigraph-Cactus callset instead of PAV as a ground truth, we obtained very similar results (Supplementary Figures 7 and 9).

### 2.4 Haplotype reconstruction of the 1kGP samples

We genotyped all 36,483,743 SNPs, 10,512,438 indels and 901,029 SV alleles contained in the HPRC2 pangenome graph across all 3,202 samples part of the 1kGP cohort and the HPRC panel samples (3,208 samples in total) [6, 36, 37] using PanGenie v4 based on Illumina data. We applied a filtering approach [3, 6, 7] (Supplementary Material), to determine a subset of reliably genotypable variants, containing 92%, 93% and 91% of all SV deletion, SV insertion and other SV alleles, respectively. Comparing the allele frequencies of these variants in the assemblies and across all 1kGP samples after genotyping, we observed high correlation (0.997 SV deletions, 0.994 SV insertions, 0.982 other SV alleles), indicating high genotyping performance (Figure 3a, Supplementary Figure 10). Additionally, allele frequencies and heterozygosities follow the Hardy Weinberg equilibrium (Figure 3b, Supplementary Figure 10). We furthermore compared our genotyped set to our previously generated 1kGP PanGenie genotypes computed based on the HGSVC3 samples [7] as well as to a short-read based SV callset generated with traditional short-read SV callers for the 1kGP cohort [36]. Since the latter set is GRCh38-based making a one-to-one comparison of SVs challenging, we instead compared the callsets based on the number of SVs genotyped as “present” (heterozygous or homozygous) per sample. The median number of SVs per sample is 27,405 for the HPRC2 based genotypes, 24,669 for the HGSVC3 genotypes and 9,261 for the Illumina-based calls (Figure 3c). When restricting to rare variants (allele frequency *<* 1%), we observed that with the HPRC2 graph, we can access hundreds of additional SVs per sample compared to the HGSVC3 genotypes and the Illumina based calls. We found medians of 1,006, 654 and 227 rare SVs per sample in HPRC, HGSVC3 and Illumina-based sets, respectively. Restricting to samples from the African continental group, we saw a median of 2,002, 1,477 and 468 of rare SVs per sample. These gains are likely a result of the larger number of assembled haplotypes in the HPRC graph, better assembly qualities compared to the HGSVC3 data and improved genotyping performance of the new PanGenie version. In particular, this shows the ability of the sampling strategy to retain relevant rare SV alleles during the haplotype sampling step.

**Fig. 3.**
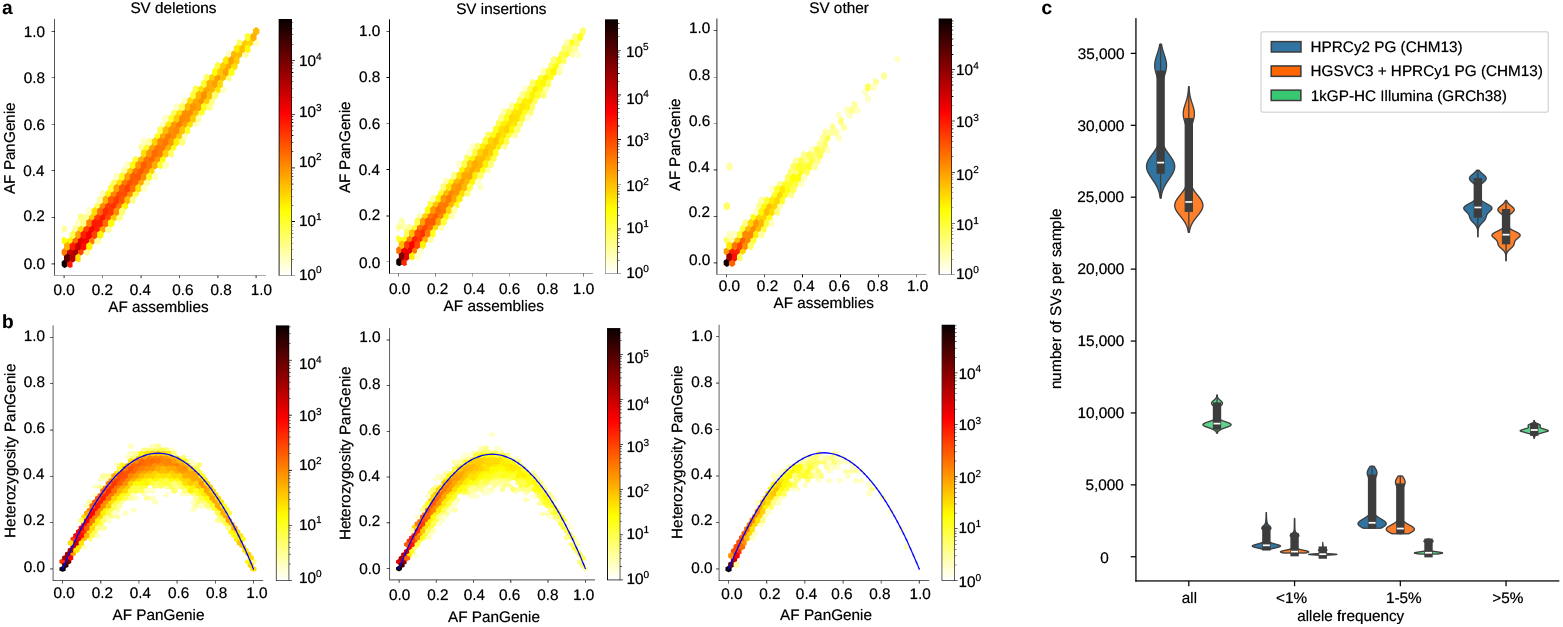
Genotyping the 1kGP cohort. **a** Comparison of allele frequencies of filtered PanGenie genotypes across 2,596 unrelated 1kGP samples and allele frequencies of the same variants in the HPRC assemblies. **b** Comparison of heterozygosities of filtered PanGenie genotypes across 2,596 unrelated 1kGP samples and PanGenie allele frequencies. **c** Number of SVs called per sample across the 1kGP cohort in the PanGenie genotypes computed based on the HPRC2 graph (blue), the PanGenie genotypes previously generated for the HGSVC3 graph (orange) and SV calls generated by traditional alignment-based methods (green).

#### 2.4.1 Creating polished consensus haplotypes

We phased our 1kGP genotypes using SHAPEIT5 [22]. We implanted phased variants back into the CHM13 reference genome to reconstruct each of the 6,416 individual haplotype sequences. Re-genotyping methods are unable to discover novel variation not represented in the pangenome graph that was used for genotyping. As a consequence, our reconstructed haplotype sequences miss (rare) variants not captured by the graph. Therefore, we developed a new polishing pipeline in order to add missed variants to our haplotype sequences.

We use short reads to call SNP and small indels with DeepVariant [38] relative to each consensus haplotype to be polished, and long reads to phase the resulting calls with WhatsHap [39] (Figure 4a). Homozygous calls mainly include variants initially missing from the graph or such that were wrongly genotyped. Heterozygous calls include mainly variants from the respective other haplotype (as reads are not separated by haplotype) and missed variants from the haplotype to be polished. We are interested in determining the latter set of calls. We can use the phasing information in order to identify them, as we would expect most of these calls to be grouped together on the haplotype reported with otherwise mostly reference alleles (Methods, Figure 4b). Furthermore, phasing enables us to split the long reads by haplotype. In this way, we call haplotype specific SVs relative to the consensus haplotype to be polished using Sniffles2 [40], providing potential SVs missed from the graph. Finally, we correct our consensus haplotypes by implanting detected SNPs, indels and SVs.

**Fig. 4.**
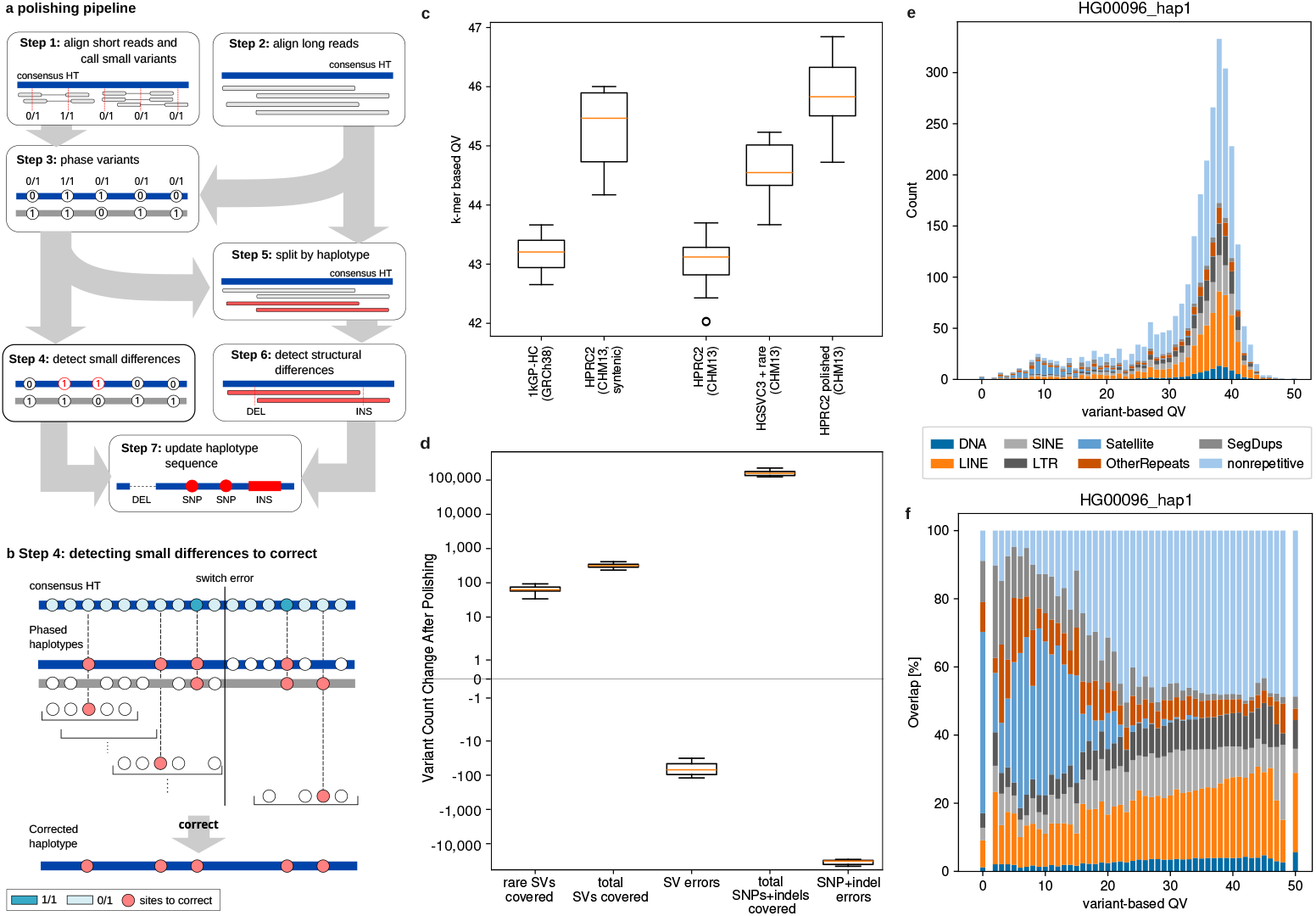
Polishing. **a** Short and long reads are aligned to the haplotype (Steps 1+2). Variants are called from short reads and phased with long reads (Step 3). A simple algorithm (see panel b) distinguishes true haplotype differences from variants originating from the opposite haplotype (Step 4). Long reads are split by haplotype based on phasing results (Step 5) and haplotype-specific SVs are called (Step 6). Small and structural variants are incorporated into the haplotype (Step 7). **b** Window-based algorithm to separate heterozygous variants arising from true differences between reads and haplotypes, and calls from the opposite haplotype. In each step, we check if the phasing of a heterozygous variant in the center of a window differs from the remaining variants. If yes, the variant is considered for polishing. Homozygous calls are always considered. **c** K-mer based QV estimates computed with Merqury for consensus haplotypes generated from phased short-read based 1kGP calls (“1kGP-HC”), from the HGSVC3 graph with PanGenie genotypes and additional rare Illumina-based SNP and indel calls (“HGSVC3 + rare”), as well as our HPRC2 unpolished and polished consensus haplotypes. Since the 1kGP haplotypes are GRCh38-based, we additionally compared results in regions shared between T2T-CHM13 and GRCh38 references (“syntenic”). Evaluation is based on 10 individuals not overlapping with HGSVC3 nor HPRC2 samples. **d** Variant count changes after polishing the PanGenie based consensus haplotypes. **c** Variant-based QVs computed across windows of length 1 Mbp along a polished consensus haplotype of HG00096. Bars are colored by the fraction of overlaps with repeat annotations and segmental duplications. **e** Histogram shown in panel c, but with each bar normalized by its height.

We evaluated our polishing pipeline based on different long read data types (PacBio HiFi and ONT) and investigated the impact of different variant types considered (SNPs, indels, SVs) on the polishing result. We computed k-mer based QV estimates with Merqury [41] to evaluate the accuracy of the polished haplotypes based on Illumina data. We also computed variant-based QV estimates based on comparison to ground truth assemblies which additionally takes structural sequence correctness into account [7]. We observed that polishing results improved with HiFi data over ONT (variant-based QV increases by around 2.5%) and that using indels and SVs for polishing led to more accurate results than polishing based on SNPs alone (variant-based QVs increase by around 2%, k-mer based QVs by around 0.5%; Supplementary Figure 11).

#### 2.4.2 Polishing increases accuracy of consensus haplotypes

We applied our polishing pipeline to a subset of 967 1kGP samples (1,934 haplotypes) for which low-coverage ONT data is available [42]. We compared our consensus haplotypes before and after polishing to consensus haplotypes generated from phased variant calls of the 1kGP cohort called from high-coverage short-read data [36] (“1kGP-HC”) as well as to consensus haplotypes we had previously generated with the HGSVC3 graph [7] (“HGSVC3 + rare”). Apart from the different graph and PanGenie version, the pipeline for consensus haplotype generation used in this manuscript differs from the HGSVC3 pipeline in the strategy used to add (rare) variation missed from the graph to the final haplotype sequences. For the HGSVC3 haplotypes, rare SNP and indel calls obtained from Illumina data for all 3,202 1kGP individuals were added to the set of graph-derived calls before phasing with SHAPEIT [7]. In this manuscript, missed (rare) variants are added through the polishing step, allowing us to additionally include SVs. To compare results, we selected 10 samples (20 haplotypes) which were neither part of the HPRC2 nor the HGSVC3 set. We computed k-mer based QV estimates using Merqury [41] based on Illumina data. Since the 1kGP-HC haplotypes are GRCh38-based, while our HPRC2 consensus haplotypes are CHM13-based, we restricted evaluation of our unpolished haplotypes to regions shared between T2T-CHM13 and GRCh38 (“syntenic”) for a fair comparison. We observed a median QV of 43.2 for the 1kGP-HC haplotypes and a median QV of 45.5 for the HPRC2 unpolished haplotypes in syntenic regions (Figure 4c), indicating that our haplotypes are more accurate. We furthermore compared our HPRC2 to HGSVC3 consensus haplotypes across all regions (both are CHM13-based). Our unpolished HPRC2 haplotypes are less accurate than the HGSVC3 haplotypes (median QVs of 43.1 and 44.5), which is explained by the additional rare variants added to the HGSVC3 haplotypes. After polishing however, our HPRC2 haplotypes reach a median QV of 45.8 (corresponding to a reduction in error rate by 25.9%), showing higher accuracy than the HGSVC3 haplotypes (Figure 4c). We also investigated the effects of a second round of polishing and observed that it can increase accuracy further (Supplementary Figure 11).

#### 2.4.3 Polishing enables capturing additional variants

In order to further compare our unpolished and polished HPRC haplotypes, we conducted an experiment in which we jointly called variants across (i) the two unpolished haplotype sequences, (ii) the two polished haplotype sequences and (iii) the two ground truth assemblies of the same sample using PAV [7]. We selected 8 individuals (16 haplotypes) which are not part of the HPRC2 graph and for which independent high quality haplotype-resolved assemblies are available from the HGSVC3 [7]. We defined three sets of variants: *rare variants* contain all ground truth SVs called in the assemblies that are not overlapping any SV in the HPRC graph, *Ground truth variants* are all variants called across the two assembly haplotypes, and *Error variants* are all variants called in the consensus haplotypes that are not called in the assembly haplotypes. We then checked how variant counts change between unpolished and polished haplotypes (Figure 4d). We found that across the 16 haplotypes evaluated, the median number of additional rare SVs captured after polishing was 63, and the median number of total ground truth SVs added was 317. At the same time, the median decrease of erroneous SVs in the polished haplotypes was 70. For SNPs and indels, the median of additional ground truth variants captured after polishing was 158,205; while the number of errors decreased by 32,199 (Figure 4d). This analysis indicates that polishing increases the accuracy of the haplotypes and indeed helps capturing (rare) variation previously missed.

#### 2.4.4 Repetitive regions are difficult to polish

We additionally checked which regions are easy and which are more difficult to polish. We used our previously developed pipeline [7] producing variant-based QV estimates for windows of 1 Mbp in length along the polished consensus haplotypes based on comparison to ground truth HGSVC3 assemblies. Since this provides local QV estimates which additionally take structural sequence correctness into account, values tend to be smaller than the k-mer based QVs computed with Merqury. This allows us to determine which regions perform well and which ones are less accurate. We plotted histograms of variant-based QVs for consensus haplotypes computed from the 1kGP-HC and HGSVC3, as well as our unpolished and polished HPRC2 haplotypes for sample HG03009 showing an increase in high QV windows for (polished) HPRC haplotypes (Supplementary Figure 12). Across all samples evaluated, we observed median variant-based QV estimates of 26.40, 33.79, 34.97 and 35.95 for the 1kGP, HGSVC3, unpolished HPRC2 and polished HPRC2 haplotypes (Supplementary Figure 13).

We additionally annotated repeats and segmental duplications in the consensus haplotypes of the evaluation samples using BISER [43] and RepeatMasker [44]. We plotted histograms of window-based QV estimates for HG00096 and analyzed overlaps of each window with these annotations (Figure 4e and 4f). Results indicate that regions with low QVs tend to have larger overlap with repetitive regions (especially Satellites and SINEs). Variant calls generated from the pangenome graph are likely less accurate in repeat associated regions and thus genotypes generated for such variants are often less accurate. Additionally, our polishing pipeline relies on alignments of short and long reads to our consensus haplotypes. Read alignment in such regions is challenging, especially for short reads. These are likely explanations for the lower accuracies of the polished haplotypes in regions overlapping repeats or segmental duplications.

### 2.5 Haplotype availability analysis

While the 462 HPRC2 haplotypes well capture common variation, we found that combining them with our set of polished haplotypes helps to better capture human diversity across highly complex and polymorphic loci of the genome. Examining 30,092 local haplotypes, extracted from 61 HGSVC3 samples over 248 non-duplicated challenging medically relevant genes [45], we observed that 41.5% of the haplotypes were completely identical to one of the haplotypes from either HPRC2 pangenome or the polished haplotypes. This represents 16.7% improvement compared to 35.6% identical haplotypes, found only in the HPRC2 assemblies (Figure 5a). Furthermore, we observed overall smaller sequence divergence between HGSVC3 haplotypes and the closest haplotype available in the extended HPRC2 + polished haplotypes panel, with 66.2% haplotypes reaching QV 40 compared to 58.4% in the original HPRC2 panel; as well as 82.1% and 89.9% haplotypes with QV ≥35 and ≥30 in the extended panel compared to 77.0% and 87.6% in the HPRC2 panel, respectively (Figure 5b).

**Fig. 5.**
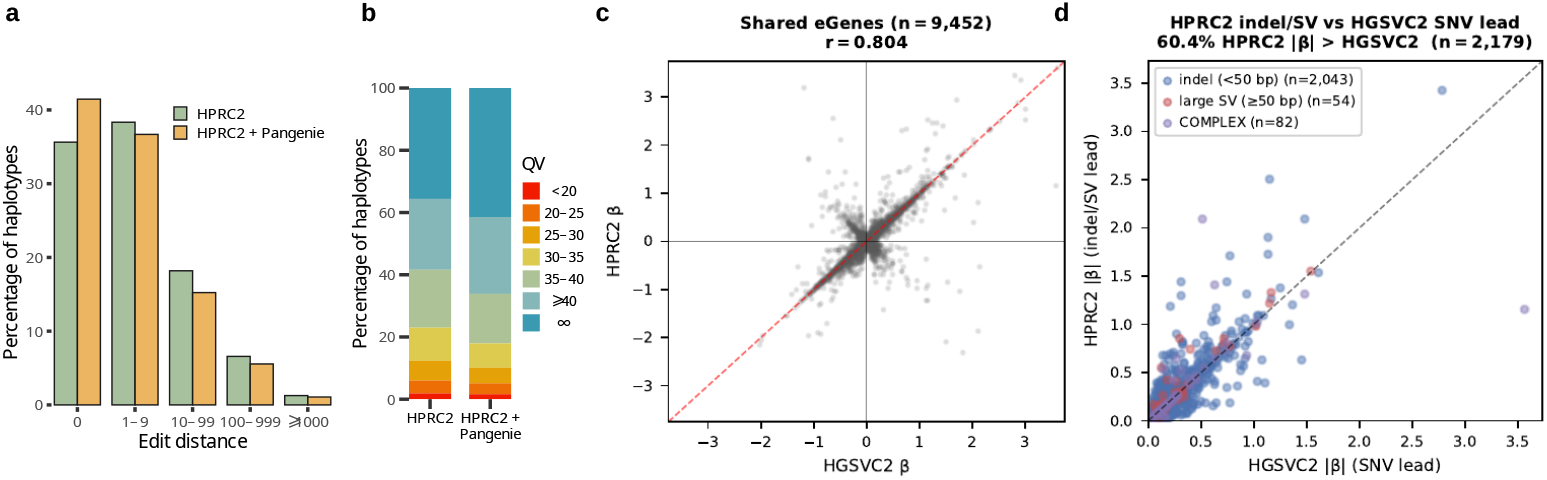
Applications. **a**–**b** Haplotype availability of the HPRC2 assemblies as well as HPRC2, enriched by the polished PanGenie haplotypes. Availability is measured as sequence divergence (edit distance in **a** and QV in **b**) from a local HGSVC3 haplotype to the closest haplotype found in the two haplotype sets, measured across 248 challenging medically relevant genes and 61 independent HGSVC3 diploid assemblies. **c** Signed effect-size concordance between HPRC2 and HGSVC2 for 9,452 eGenes detected in both panels at FDR 5% (Pearson *r* = 0.804). Each point is one shared eGene; dashed line is the diagonal. **d** Among shared eGenes where HPRC2 assigns an indel or SV as the lead variant and HGSVC2 assigns an SNP (*n* = 2,179), HPRC2 shows the larger 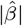 in 60.4% of cases versus 40.0% in a matched control where both panels assign an SNP lead (*n* = 6,444; *χ*^2^ = 273, *p <* 10^*−*60^). Colors indicate HPRC2 lead variant category.

### 2.6 eQTL analysis

To demonstrate how the HPRC2 panel supports population-scale functional genomics, we performed cis-eQTL mapping in 397 Geuvadis lymphoblastoid cell line samples, 20 of which contributed haplotypes to the HPRC2 panel, extending the analysis of Ebert et al. [3]. Each sample was genotyped with PanGenie v4.2.1 against the HPRC2 graph and the HGSVC2 PAV Freeze4 panel used in Ebert et al.; cis-eQTLs were mapped with tensorQTL [46] (MAF ≥ 0.01, 60 expression PCs, *±*1 Mb cis-window, FDR 5%).

Both panels exceed the Ebert 2021 benchmark of 9,401 eGenes, identifying 10,222 (HPRC2) and 10,456 (HGSVC2) eGenes (92.5% and 90.4% mutual detection). Across 9,452 shared eGenes, lead-variant effect sizes are closely correlated (*r* = 0.804; Figure 5c), indicating both panels recover the same common-variant cis-regulatory architecture.

HPRC2 assigns a non-SNV variant (insertion, deletion, or multi-nucleotide variant) as the lead at more eGenes than HGSVC2 (3,200 versus 978). HPRC2 lead variants have a larger median absolute effect size overall (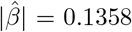 versus 0.1257 for HGSVC2). Where the panels disagree on lead variant type – HPRC2 finds an indel or SV lead, HGSVC2 an SNV (*n* = 2,097, excluding 82 HPRC2-only multi-nucleotide variant leads absent from the PAV callset) – the HPRC2 lead has the larger 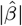 in 61.0% of cases; in a control where both panels assign an SNV lead (*n* = 6,444), only 40.0% (*χ*^2^ = 283, *p <* 10^*−*62^; Figure 5d), indicating the effect-size advantage is concentrated at loci where HPRC2 offers an indel or SV lead rather than reflecting a uniform panel-level effect. The sample size (*n* = 397) limits power for rare-variant and population-stratified analyses, left to future work in larger cohorts.

## 3 Discussion

Pangenome graphs constructed from highly accurate haplotype resolved assemblies enable downstream analyses of genomes based on short read data. We have presented a new version of our previously introduced genotyping software PanGenie, which now includes a new haplotype sampling step that reduces the number of input haplotypes represented in the graph to speed up the subsequent genotyping step. This allows PanGenie to be applied to pangenome graphs exceeding 254 haplotypes, which previous versions were limited to. We showed that PanGenie v4 is 12*×* faster (5*×* faster in terms of walltime using 24 cores) and more memory efficient than the previous version, while being more accurate than the other state-of-the-art method KAGE. We applied PanGenie to the 1kGP cohort and phased resulting genotypes with SHAPEIT5, enabling the construction of consensus haplotype sequences for all 3,202 samples. We further polished haplotypes for a subset of 967 samples for which low coverage ONT data was available, which increased median QVs from 43 (unpolished) to 46 (polished).

We demonstrate that PanGenie enables fast eQTL analysis based on the HPRC2 pangenome variants, capturing the same core cis-regulatory architecture as earlier panels, while revealing more indel and structural variant-associated eQTLs. This highlighting its potential to improve the discovery of functional genetic variation beyond SVs.

We furthermore showed that our polished haplotypes better capture human diversity than the HPRC2 assemblies alone for highly complex and polymorphic loci of the genome, providing an improved and more complete pangenome reference that downstream applications can benefit from.

However, there are still limitations. While we expect PanGenie v4 to be able to handle graphs with up to a few thousands of haplotypes, an issue could be memory usage. Current file formats like the GFA format used to store these graphs, as well as the VCF format used to store variants derived from the graph will likely become very large. This will increase memory consumption of PanGenie and other downstream analysis tools that rely on these files. Even for the HPRC2 data, file size of the gzip-compressed VCF and GFA files reached 6GB and 60GB, respectively. These issues could be addressed by introducing new file formats that better scale to large graphs, as well as implementing more space-efficient data structures to store variant and haplotype information in PanGenie.

As we were unable to get KAGE2 (v2.0.0) to run on our data, we used a slightly modified version of KAGE for our benchmarking experiments which was previously developed by the *All of Us* consortium [35]. Although changes were relatively minor and mainly affected scalability and robustness, performance regressions might have inadvertently been introduced.

For polishing, we rely on short and long read data of a certain sample to correct residual differences in the consensus haplotypes. The bottlenecks of our current polishing pipeline are the alignment of short and long reads to the haplotype, as well as the small variant calling with DeepVariant, which combined take up more than 90% of the runtime. Here, we will explore different options to speed up these steps, including experimenting with alternative variant callers.

In conclusion, we showed that using a pangenome graph in combination with short read data, enables accurate reconstruction of haplotype sequences of human individuals. Including low coverage ONT data helps removing residual errors and add missed variation to these consensus haplotypes, producing a pangenome resource able to capture diversity at complex genomic regions.

## 4 Online Methods

### 4.1 PanGenie v4

We developed a new version of PanGenie, PanGenie v4, which includes an additional haplotype sampling step prior to genotyping. The aim is to reduce the full set of haplotypes to a smaller set, tailored to the sample to be genotyped, in order to speed up the subsequent genotyping step. During the haplotype sampling step, we construct new haplotype sequences which are mosaics of the original haplotypes in the graph. These haplotypes are constructed in a sample-specific manner by prioritizing path fragments more similar to the individuals genome sequence, measured in terms of shared unique k-mer sequences between haplotype paths and sequencing reads of a sample.

#### 4.1.1 Identifying unique k-mers

For each bubble, we compute a set of k-mers that are unique to the allele sequences represented in the bubble. In contrast to how we had defined unique k-mers in the original PanGenie model, we additionally require the k-mers to occur in exactly one allele, i.e. we no longer include k-mers unique to a bubble region but shared between alleles within the same bubble. We count k-mers in the sequencing reads of the sample to be genotyped to determine counts for all unique k-mers, as well as the average k-mer coverage [9]. We define the total set of unique k-mers as *K* and *Count*(*k*) denotes the count of a unique k-mer *k ∈ K* in the sequencing data of the sample.

#### 4.1.2 Haplotype sampling algorithm

Our haplotype sampling algorithm is iterative. In each iteration, we construct a new haplotype sequence based on a dynamic programming (DP) algorithm.

Given *M* bubbles and *N* haplotypes in the input pangenome graph, the DP table consists of *M* columns and *N* rows. Each bubble *j* of the graph contains up to *N* different allele sequences, which we refer to as *a*_*jk*_ in the following (Supplementary Figure 1). We furthermore define a mapping between haplotypes and the alleles they cover in each bubble as *Allele*_*j*_(*h*_*i*_). For each allele *a*_*jk*_ in bubble *j*, we denote the set of unique k-mers it contains as *Kmers*_*j*_(*a*_*jk*_) = {*k* ∈ *K*|*k* ∈ *a*_*jk*_}. Additionally, we define the set of read k-mers of each allele as *PresentKmers*_*j*_(*a*_*jk*_) = {*k* ∈ *K*|*k* ∈ *a*_*jk*_ ∧ *Count*(*k*) ≥ 3}. Here, we assume a k-mer to be present in the sequencing data of the sample if it was seen at least 3 times.

##### Local Costs

We refer to the entries of the DP table as *D*_*i,j*_ and each such cell is associated with a local cost which we refer to as *L*(*D*_*i,j*_). We assign a cost to each allele of a bubble *j* based on how many of the unique k-mers it carries are also present in the sequencing data of the sample. These costs are defined as:

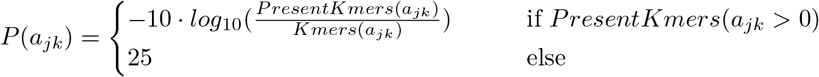

The local costs of each cell *D*_*i,j*_ corresponding to a haplotype *h*_*i*_ are then defined by the allele it carries:

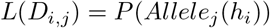

##### Transition Costs

We denote transition costs between adjacent columns *j −* 1 and *j* as *T* (*D*_*i,j*_ | *D*_*x,j−*1_). Transition costs are computed based on the Li-Stephens model [32] and are defined similar to how they are used in PanGenie [9]. Given a recombination rate *r*, an effective population size *N*_*e*_ and the distance *t* (in basepairs) between two ascending bubbles *j −* 1 and *j*, we define:

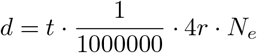

The Li-Stephens transition probability is computed as:

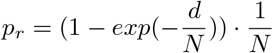

We use log-scaled transition probabilities to define costs of switching from one haplotype to a different one between two adjacent columns:

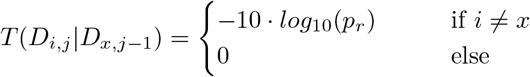

###### Algorithm

In each iteration of the sampling algorithm, one haplotype path is constructed by finding the path with minimal costs through the DP table. The index of the respective cell in each column is removed for the next iteration of the DP algorithm, i.e. each haplotype can only be selected once per column. We define a set *S*_*j*_ containing indices of all haplotypes not yet selected in column *j* for all *j* ∈ {1, …, *M*}. We run *R* iterations of our DP algorithm in order to construct *R* haplotypes. After each iteration, we update the local costs of all alleles that are covered by the selected haplotype path by adding a penalty score. This encourages the algorithm to select other alleles in the following steps to achieve a larger spectrum of possible alleles covered.

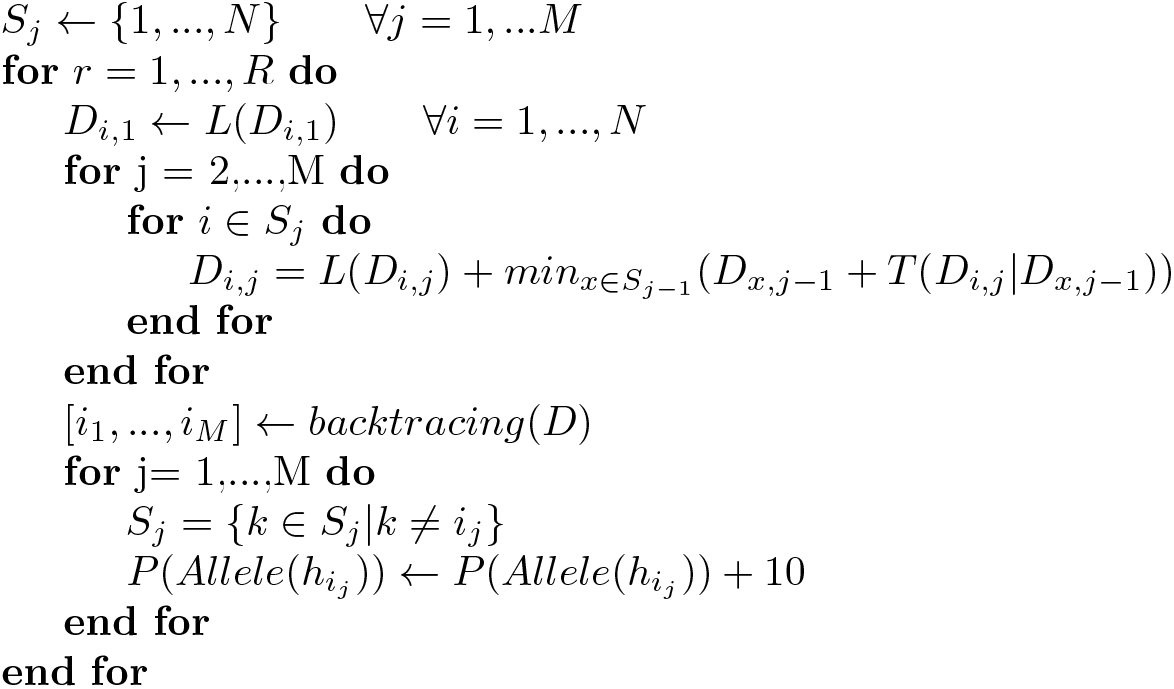

The function *backtracing*(*D*) performs backtracing on the DP matrix *D* starting from the smallest cell in the last column *D*_*i,M*_ in order to determine the path through all columns with the lowest cost. It returns a list of indices indicating which haplotype was selected in each column.

##### Asymptotic Runtime

One column of the DP matrix can be filled in *O*(*N*) time by precomputing minima from the previous column (which can also be done in *O*(*N*) time). One run of the DP algorithm can thus be performed in *O*(*M · N*) time. With *R* iterations, the total runtime of the haplotype sampling algorithm is thus *O*(*R · M · N*).

### 4.2 Preprocessing of the HPRC2 graph

We used the CHM13-based HPRC v2.0 graph generated with Minigraph-Cactus [8]. Besides a GFA file, the Minigraph-Cactus pipeline produces a VCF file encoding the top-level bubbles of the graph with respective genotypes for all 231 graph samples. We performed some preprocessing on this VCF. First, we converted genotypes of all male samples on chrX and chrY to a homozygous representation by duplicating the haplotype carrying less missing alleles (“.”) in the VCF. Next, we removed all records for which more than 20% of all haplotypes carried a missing allele (“.”) and postprocessed the remaining bubbles using a decomposition pipeline that we had introduced earlier [6, 7]. This pipeline decomposes the bubbles based on the allele traversals of the haplotypes to detect variant alleles nested inside of the bubbles [6]. It adds annotations to the VCF encoding these nested variants in the ID field (“bubble VCF”), and generates an additional VCF reporting each nested alleles as a separate, biallelic record (“callset VCF”). We then use the “bubble VCF” as a input to PanGenie to genotype bubbles and use the “callset VCF” to convert the bubble genotypes to genotypes of all nested variant alleles [6].

### 4.3 Running PanGenie on HPRC data

We first run indexing once on the “bubble VCF” using the command: `PanGenie-index -v <bubble VCF> -r <reference FASTA> -o index -t 24`. For each sample, we then compute genotypes using: `PanGenie -f index -i <sample> reads.fasta -o pangenie <sample> -t 24 -j 24 -s <sample>`. Finally, we convert bubble genotypes to genotypes for variant alleles (after decomposition): `cat pangenie <sample> genotyping.vcf | python3 convert-to-biallelic.py <callset VCF> | bgzip > <sample> genotyping bi.vcf.gz`

### 4.4 Polishing algorithm

Our polishing pipeline requires the haplotype sequence to be provided in FASTA format as well as short read and long read sequencing data for the respective individual. First, we align the short reads to the haplotype sequence using strobealign [47] and call SNPs and indels with DeepVariant [38] (Figure 4a, Step 1). We align the long reads with minimap2 [48] (Figure 4a, Step 2). Next, we use the long read alignments to phase the variant calls with WhatsHap [39] (Figure 4a, Step 3). Both sequencing datasets contain reads from both haplotypes of an individual and not just the reads corresponding to the haplotype that we aim to polish. Therefore, heterozygous calls include mainly variants from the respective other haplotype, which should ideally be grouped together after phasing, and missed variants from the haplotype to be polished, ideally located on the opposite phased block with otherwise mostly reference alleles. For polishing, we are interested in determining the latter set of calls, as they represent sites in the haplotype sequence that we want to polish. In order to determine these calls (Figure 4a, Step 4), we use a simple algorithm (Figure 4b) which slides a window along the heterozygous phased variants and in each step, checks whether the variant in the center is phased differently than the majority of the other variants included in the window. If this is the case, we mark this variant as a position to be polished. Then, we slide the window by one position and repeat the process until we have covered the whole haplotype sequence. All SNP and indel calls detected in this way represent positions to be corrected in the haplotype to be polished (referred to *corr*_*a*_ in the following). Additionally, all homozygous SNP and indel calls reported by DeepVariant need to be corrected (*corr*_*b*_). The default number of heterozygous variants considered per window is set to 25.

So far, our polishing pipeline has only considered SNPs and indels. In order to incorporate SVs, we split the long reads by haplotype in order to only keep the set of reads corresponding to the haplotype that we want to polish (Figure 4a, Step 5). We do this based on the phasing results using WhatsHap’s `haplotag` and `split` commands. We then call haplotype specific SVs relative to the haplotype to be polished using Sniffles2 [40] (*corr*_*c*_, Figure 4a, Step 6).

In the last step, we polish the haplotype by incorporating all detected heterozygous small variant positions (*corr*_*a*_), all homozygous small variant positions (*corr*_*b*_) and all called SVs (*corr*_*c*_) into the sequence using `bcftools consensus` [49] (Figure 4a, Step 7). The output of the polishing pipeline is a FASTA file containing the polished haplotype sequence.

## Supplementary information

### Data Availability

Genotypes across all 1kGP cohort samples as well as consensus haplotypes before and after polishing are available at: https://s3-us-west-2.amazonaws.com/human-pangenomics/index.html?prefix=pangenomes/scratch/2026_03_30_pangenie/

### Code Availability

The haplotype sampling algorithm is implemented as part of the PanGenie software which is available at: https://github.com/eblerjana/pangenie. Our polishing pipeline is available at: https://github.com/eblerjana/polishing-pipeline. Our pipelines used to run the experiments for this manuscript are available at: https://github.com/eblerjana/hprc2-companion

### Author Contributions

JE and TM designed the haplotype sampling algorithm, the polishing pipeline and the overall study. JE implemented the haplotype sampling algorithm, the polishing pipeline and the pipelines to genotype and polish the 1kGP cohort as well as the benchmarking experiments. PE ran RepeatMasker on the polished consensus haplotypes. SKL produced all KAGE results. TP conducted the haplotype availability experiments. AB and BP produced eQTL results. JE, SKL, TP, AB and TM wrote the manuscript and all authors contributed edits and comments.

## Acknowledgements

Computational infrastructure and support were provided by the Centre for Information and Media Technology at Heinrich Heine University Düsseldorf. We would like to acknowledge the National Human Genome Research Institute (NHGRI) for funding the following grants supporting the creation of the human pangenome reference: U41HG010972, U01HG010971, U01HG013760, U01HG013755, U01HG013748, U01HG013744, R01HG011274, and the Human Pangenome Reference Consortium (BioProject ID: PRJNA730823).

## Human Pangenome Reference Consortium

Derek Albracht^1^, Ivan A. Alexandrov^2^, Jamie Allen^3^, Alawi A. Alsheikh-Ali^4^, Nicolas Altemose^5^, Casey Andrews^6^, Dmitry Antipov^7^, Lucinda Antonacci-Fulton^1^, Mobin Asri^8^, Marcelo Ayllon^9^, Jennifer R. Balacco^10^, Floris P. Barthel^11^, Edward A. Belter Jr^1^, Halle D. Bender^8^, Andrew P. Blair^8^, Davide Bolognini^12^, Katherine E. Bonini^13^, Christina Boucher^14^, Guillaume Bourque^15,16,17^, Silvia Buonaiuto^18^, Shuo Cao^18^, Andrew Carroll^19^, Ann M. Mc Cartney^8^, Monika Cechova^8^, Mark J.P. Chaisson^20^, Pi-Chuan Chang^19^, Xian Chang^8^, Jitender Cheema^3^, Haoyu Cheng^21^, Claudio Ciofi^22^, Hiram Clawson^8^, Sarah Cody^1^, Vincenza Colonna^18^, Holland C. Conwell^23^, Robert Cook-Deegan^24^, Mark Diekhans^8^, Maria Angela Diroma^22^, Daniel Doerr^25,26,27^, Zheng Dong^6^, Danilo Dubocanin^5^, Richard Durbin^28,29^, Jana Ebler^25,30^, Evan E. Eichler^9,31^, Jordan M. Eizenga^8^, Parsa Eskandar^8^, Eddie Ferro^14^, Anna-Sophie Fiston-Lavier^32,33^, Sarah M. Ford^23^, Willard W. Ford^34^, Giulio Formenti^10^, Adam Frankish^3^, Mallory A. Freeberg^3^, Qichen Fu^6^, Stephanie M. Fullerton^35^, Robert S. Fulton^1^, Yan Gao^36^, Gage H. Garcia^9^, Obed A. Garcia^37^, Joshua M.V. Gardner^8^, Shilpa Garg^38^, Erik Garrison^18^, Nanibaa’ A. Garrison^39,40,41^, John E. Garza^1^, Margarita Geleta^42^, Mohammadmersad Ghorbani^43^, Tina A. Graves-Lindsay^1^, Richard E. Green^23^, Cristian Groza^44^, Bida Gu^20^, Andrea Guarracino^11,18^, Melissa Gymrek^45^, Maximilian Haeussler^8^, Leanne Haggerty^3^, Ira M. Hall^46,47^, Nancy F. Hansen^7^, Yue Hao^11^, Mohammad Amiruddin Hashmi^4^, David Haussler^8^, Prajna Hebbar^8^, Peter Heringer^25,26,27^, Glenn Hickey^8^, Todd L. Hillaker^8^, S. Nakib Hossain^3^, Neng Huang^36,48^, Sarah E. Hunt^3^, Toby Hunt^3^, Alexander G. Ioannidis^5,8^, Nafiseh Jafarzadeh^8^, Nivesh Jain^10^, Erich D. Jarvis^10,31^, Maryam Jehangir^11^, Juan Jiang^6^, Eimear E. Kenny^13^, Juhyun Kim^7^, Bonhwang Koo^10^, Sergey Koren^7^, Milinn Kremitzki^1,6^, Charles H. Langley^49^, Ben Langmead^50^, Heather A. Lawson^6^, Daofeng Li^6^, Heng Li^36,48^, Wen-Wei Liao^46,47^, Jiadong Lin^9^, Tianjie Liu^6^, Glennis A. Logsdon^51^, Ryan Lorig-Roach^8^, Jonathan LoTempio Jr^52^, Hailey Loucks^8^, Jane E. Loveland^3^, Jianguo Lu^53^, Shuangjia Lu^46,47^, Julian K. Lucas^8^, Walfred Ma^20^, Juan F. Macias-Velasco^1,6,54^, Kateryna D. Makova^55^, Maximillian G. Marin^36,48^, Christopher Markovic^1^, Tobias Marschall^25,30^, Franco L. Marsico^18^, Fergal J. Martin^3^, Mira Mastoras^8^, Capucine Mayoud^32^, Brandy McNulty^8^, Jack A. Medico^10^, Julian M. Menendez^8^, Karen H. Miga^8^, Anna Minkina^56^, Matthew W. Mitchell^57^, Saswat K. Mohanty^58^, Younes Mokrab^43,59,60^, Jean Monlong^61^, Shabir Moosa^43^, Avelina Moreno-Ochando^62,63^, Shinichi Morishita^64^, Jonathan M. Mudge^3^, Katherine M. Munson^9^, Njagi Mwaniki^65^, Nasna Nassir^4^, Chiara Natali^22^, Shloka Negi^8^, Lingbin Ni^9^, Adam M. Novak^8^, Faith Okamoto^8^, Pilar N. Ossorio^66^, Chie Owa^64^, Sadye Paez^10^, Benedict Paten^8^, Clelia Peano^12,67^, Adam M. Phillippy^7^, Brandon D. Pickett^7^, Laura Pignata^18^, Nadia Pisanti^65^, David Porubsky^9,68^, Pjotr Prins^18^, Anandi Radhakrishnan^8^, T. Rhyker Ranallo-Benavidez^11^, Brian J. Raney^8^, Mikko Rautiainen^69^, Alessandro Raveane^12^, Luyao Ren^9,31^, Arang Rhie^7^, Fedor Ryabov^70,71^, Samuel Sacco^23^, Farnaz Salehi^18^, Michael C. Schatz^50,72^, Laura B. Scheinfeldt^73^, Aarushi Sehgal^34^, William E. Seligmann^23^, Mahsa Shabani^74^, Kishwar Shafin^19^, Shadi Shahatit^32^, Ruhollah Shemirani^13^, Vikram S. Shivakumar^50^, Swati Sinha^3^, Jouni Sirén^8^, Linnéa Smeds^58^, Steven J. Solar^7^, Marco Sollitto^10,22^, Nicole Soranzo^12,28,75^, Andrew B. Stergachis^9,56^, Marie-Marthe Suner^3^, Yoshihiko Suzuki^64^, Arda Söylev^25,30^, Ahmad Abou Tayoun^76,77^, Jack A.S. Tierney^3^, Chad Tomlinson^1^, Francesca Floriana Tricomi^3^, Mohammed Uddin^4,78^, Matteo Tommaso Ungaro^23,79^, Rahul Varki^14^, Flavia Villani^18^, Ivo Violich^8^, Mitchell R. Vollger,^56^, Brian P. Walenz^7^, Charles Wang^80^, Lisa E. Wang^13^, Ting Wang^1,6,54^, Aaron M. Wenger^81^, Conor V. Whelan^10^, Zilan Xin^6^, Zheng Xu^6^, Kai Ye^82^, DongAhn Yoo^9^, Wenjin Zhang^6^, Ying Zhou^36^, Xiaoyu Zhuo^6^, Giulia Zunino^12^

^1^ McDonnell Genome Institute, Washington University School of Medicine, St. Louis, MO 63108, USA ^2^ Department of Human Molecular Genetics and Biochemistry, Faculty of Medical and Health Sciences, Tel Aviv University, Tel Aviv 69978, Israel ^3^ European Molecular Biology Laboratory, European Bioinformatics Institute (EMBL-EBI), Wellcome Genome Campus, Hinxton, Cambridge CB10 1SD, UK ^4^ Center for Applied and Translational Genomics (CATG), Mohammed Bin Rashid University of Medicine and Health Sciences, Dubai Health, Dubai, UAE ^5^ Department of Genetics, Stanford University, Palo Alto, CA 94304 USA ^6^ Department of Genetics, Washington University School of Medicine, St. Louis, MO 63110, USA ^7^ Genome Informatics Section, Center for Genomics and Data Science Research, National Human Genome Research Institute, National Institutes of Health, Bethesda, MD 20892, USA ^8^ UC Santa Cruz Genomics Institute, University of California, Santa Cruz, CA 95060, USA ^9^ Department of Genome Sciences, University of Washington School of Medicine, Seattle, WA 98195, USA ^10^ The Vertebrate Genome Laboratory, The Rockefeller University, New York, NY 10065, USA ^11^ Bioinnovation and Genome Sciences, The Translational Genomics Research Institute (TGen), Phoenix, AZ 85004, USA ^12^ Human Technopole, Milan, Italy ^13^ Institute for Genomic Health, Icahn School of Medicine at Mount Sinai, New York, NY 10029, USA ^14^ Department of Computer and Information Science and Engineering, University of Florida, Gainesville, FL 32611, USA ^15^ Canadian Center for Computational Genomics, McGill University, Montréal, QC H3A 0G1, Canada ^16^ Department of Human Genetics, McGill University, Montréal, QC H3A 0G1, Canada ^17^ Victor Phillip Dahdaleh Institute of Genomic Medicine, Montréal, QC H3A 0G1, Canada ^18^ Department of Genetics, Genomics and Informatics, University of Tennessee Health Science Center, Memphis, TN 38163, USA ^19^ Google LLC, Mountain View, CA 94043, USA ^20^ Quantitative and Computational Biology, University of Southern California, Los Angeles, CA 90089, USA ^21^ Department of Biomedical Informatics and Data Science, Yale School of Medicine, New Haven, CT 06510, USA ^22^ Department of Biology, University of Florence, Sesto Fiorentino, FI 50019, Italy ^23^ Department of Ecology and Evolutionary Biology, University of California, Santa Cruz, CA 95060, USA ^24^ Arizona State University, Consortium for Science, Policy & Outcomes, Washington, DC 20006, USA ^25^ Center for Digital Medicine, Heinrich Heine University Düsseldorf, Düsseldorf, NRW, DE ^26^ Department for Endocrinology and Diabetology at the Medical Faculty and University Hospital Düsseldorf, Heinrich Heine University Düsseldorf, Düsseldorf, NRW, DE ^27^ Paul-Langerhans-Group Computational Diabetology, German Diabetes Center (DDZ) and Leibniz Institute for Diabetes Research, Düsseldorf, NRW, DE ^28^ Wellcome Sanger Institute, Genome Campus, Hinxton, CB10 1RQ, UK ^29^ Department of Genetics, University of Cambridge, Cambridge, CB2 3EH, UK ^30^ Institute for Medical Biometry and Bioinformatics, Medical Faculty and University Hospital Düsseldorf, Heinrich Heine University, Düsseldorf, NRW, DE ^31^ Howard Hughes Medical Institute, Chevy Chase, MD 20815, USA ^32^ ISEM, Univ Montpellier, CNRS, IRD, Montpellier, FR ^33^ Institut Universitaire de France, Paris, FR ^34^ Department of Computer Science and Engineering, University of California San Diego, La Jolla, CA 92093, USA ^35^ Department of Bioethics & Humanities, University of Washington School of Medicine, Seattle, WA 98195, USA ^36^ Department of Data Science, Dana-Farber Cancer Institute, Boston, MA 02215, USA ^37^ Department of Anthropology, University of Kansas, Lawrence, KS 66045, USA ^38^ School of Health Sciences, University of Manchester, Manchester M13 9PL, UK ^39^ Traditional, ancestral and unceded territory of the Gabrielino/Tongva peoples, Institute for Society & Genetics, University of California, Los Angeles, Los Angeles, CA 90095, USA ^40^ Traditional, ancestral and unceded territory of the Gabrielino/Tongva peoples, Institute for Precision Health, David Geffen School of Medicine, University of California, Los Angeles, Los Angeles, CA 90095, USA ^41^ Traditional, ancestral and unceded territory of the Gabrielino/Tongva peoples, Division of General Internal Medicine & Health Services Research, David Geffen School of Medicine, University of California, Los Angeles, Los Angeles, CA 90095, USA ^42^ Department of Electrical Engineering and Computer Science, University of California, Berkeley, Berkeley, CA 94720, USA ^43^ Medical and Population Genomics Lab, Sidra Medicine, Doha, Qatar ^44^ Montreal Heart Institute, Montréal, QC, Canada ^45^ Department of Pediatrics, University of California San Diego, La Jolla, CA 92093, USA ^46^ Center for Genomic Health, Yale University School of Medicine, New Haven, CT 06510, USA ^47^ Department of Genetics, Yale University School of Medicine, New Haven, CT 06510, USA ^48^ Department of Biomedical Informatics, Harvard Medical School, Boston, MA 02115, USA ^49^ Department of Evolution and Ecology and the Center for Population Biology, University of California, One Shields, Davis, CA 95616, USA ^50^ Department of Computer Science, Johns Hopkins University, Baltimore, MD 21218, USA ^51^ Department of Genetics, Epigenetics Institute, Perelman School of Medicine, University of Pennsylvania, Philadelphia, PA 19104, USA ^52^ Department of Pediatrics, Division of Genetics, School of Medicine, University of California, Irvine, CA 92697, USA ^53^ Sun Yat-sen University, Guangzhou, China ^54^ Edison Family Center for Genome Sciences & Systems Biology, Washington University School of Medicine, St. Louis, MO 63110, USA ^55^ Department of Biology and Center for Medical Genomics, Penn State University, University Park, PA 16802, USA ^56^ Division of Medical Genetics, Department of Medicine, University of Washington School of Medicine, Seattle, WA 98195, USA ^57^ The Jackson Laboratory for Genomic Medicine, Farmington, CT 06032, USA ^58^ Department of Biology, Penn State University, University Park, PA 16802, USA ^59^ Department of Biomedical Science, College of Health Sciences, Qatar University, Doha, Qatar ^60^ Department of Genetic Medicine, Weill Cornell Medicine-Qatar, Doha, Qatar ^61^ IRSD - Digestive Health Research Institute, University of Toulouse, INSERM, INRAE, ENVT, UPS, Toulouse, FR ^62^ MATCH biosystems, S.L., Elche, Spain ^63^ Universidad Miguel Hernández de Elche, Elche, Spain ^64^ Department of Computational Biology and Medical Sciences, The University of Tokyo, Kashiwa, Chiba 277-8561, Japan ^65^ Department of Computer Science, University of Pisa, Pisa, Italy ^66^ Law School, University of Wisconsin-Madison, Madison, WI 53706, USA ^67^ Institute of Genetics and Biomedical Research, UoS of Milan, National Research Council, Milan, Italy ^68^ Genome Biology Unit, European Molecular Biology Laboratory (EMBL), Heidelberg, DE ^69^ Institute for Molecular Medicine Finland, Helsinki Institute of Life Science, University of Helsinki, Helsinki, Finland ^70^ The Center for Bio- and Medical Technologies, Moscow, RUS ^71^ Centre for Biomedical Research and Technology, HSE University, Moscow, RUS ^72^ Department of Biology, Johns Hopkins University, Baltimore, MD 21218, USA ^73^ Coriell Institute for Medical Research, Camden, NJ 08103, USA ^74^ University of Amsterdam, Amsterdam, Netherlands ^75^ School of Clinical Medicine, University of Cambridge, Cambridge, CB2 0SP, UK ^76^ Center for Genomic Discovery, Mohammed Bin Rashid University, Dubai Health, UAE ^77^ Dubai Health Genomic Medicine Center, Dubai Health, UAE ^78^ GenomeArc Inc, Mississauga, ON, Canada ^79^ Department of Biology and Biotechnologies “Charles Darwin”, University of Rome “La Sapienza”, Rome 00185, IT ^80^ Center for Genomics, Loma Linda University School of Medicine, Loma Linda, CA 92350, USA ^81^ PacBio, Menlo Park, CA 94025, USA ^82^ The first affiliated hospital of Xi’an Jiaotong University, Xi’an Jiaotong University, Xi’an, Shaanxi, 710049, China

## References

[1] Wenger, A. M. et al. Accurate circular consensus long-read sequencing improves variant detection and assembly of a human genome. Nature Biotechnology 37, 1155–1162 (2019).

[2] Nurk, S. et al. The complete sequence of a human genome. Science 376, 44–53 (2022).

[3] Ebert, P. et al. Haplotype-resolved diverse human genomes and integrated analysis of structural variation. Science 372, eabf7117 (2021).

[4] Cheng, H., Concepcion, G. T., Feng, X., Zhang, H. & Li, H. Haplotype-resolved de novo assembly using phased assembly graphs with hifiasm. Nature Methods 18, 170–175 (2021).

[5] Rautiainen, M. et al. Telomere-to-telomere assembly of diploid chromosomes with verkko. Nature Biotechnology 41, 1474–1482 (2023).

[6] Liao, W.-W. et al. A draft human pangenome reference. Nature 617, 312–324 (2023).

[7] Logsdon, G. A. et al. Complex genetic variation in nearly complete human genomes. Nature 644, 430–441 (2025).

[8] Hickey, G. et al. Pangenome graph construction from genome alignments with minigraph-cactus. Nature Biotechnology 42, 663–673 (2024).

[9] Ebler, J. et al. Pangenome-based genome inference allows efficient and accurate genotyping across a wide spectrum of variant classes. Nature Genetics 54, 518–525 (2022).

[10] Consortium,. G. P. et al. A global reference for human genetic variation. Nature 526, 68 (2015).

[11] Sudlow, C. et al. UK biobank: an open access resource for identifying the causes of a wide range of complex diseases of middle and old age. PLoS Medicine 12, e1001779 (2015).

[12] The All of Us Research Program Genomics Investigators. Genomic data in the All of Us Research Program. Nature 627, 340–346 (2024).

[13] Metspalu, A. The Estonian genome project. Drug Development Research 62, 97–101 (2004).

[14] Kurki, M. I. et al. FinnGen provides genetic insights from a well-phenotyped isolated population. Nature 613, 508–518 (2023).

[15] Minton, K. The FinnGen study: disease insights from a ‘bottlenecked’population. Nature Reviews Genetics 24, 207–207 (2023).

[16] Zhao, X. et al. Expectations and blind spots for structural variation detection from long-read assemblies and short-read genome sequencing technologies. The American Journal of Human Genetics 108, 919–928 (2021).

[17] Weischenfeldt, J., Symmons, O., Spitz, F. & Korbel, J. O. Phenotypic impact of genomic structural variation: insights from and for human disease. Nature Reviews Genetics 14, 125–138 (2013).

[18] Sudmant, P. H. et al. An integrated map of structural variation in 2,504 human genomes. Nature 526, 75–81 (2015).

[19] Sirén, J. et al. Pangenomics enables genotyping of known structural variants in 5202 diverse genomes. Science 374, abg8871 (2021).

[20] Grytten, I., Dagestad Rand, K. & Sandve, G. K. Kage: fast alignment-free graph-based genotyping of snps and short indels. Genome Biology 23, 209 (2022).

[21] Grytten, I., Rand, K. D. & Sandve, G. K. Kage 2: Fast and accurate genotyping of structural variation using pangenomes. bioRxiv 2023–12 (2023).

[22] Hofmeister, R. J., Ribeiro, D. M., Rubinacci, S. & Delaneau, O. Accurate rare variant phasing of whole-genome and whole-exome sequencing data in the UK Biobank. Nature Genetics 55, 1243–1249 (2023).

[23] Rubinacci, S., Ribeiro, D. M., Hofmeister, R. J. & Delaneau, O. Efficient phasing and imputation of low-coverage sequencing data using large reference panels. Nature Genetics 53, 120–126 (2021).

[24] Rubinacci, S., Hofmeister, R. J., Sousa da Mota, B. & Delaneau, O. Imputation of low-coverage sequencing data from 150,119 UK Biobank genomes. Nature Genetics 55, 1088–1090 (2023).

[25] Walker, B. J. et al. Pilon: an integrated tool for comprehensive microbial variant detection and genome assembly improvement. PloS One 9, e112963 (2014).

[26] Zimin, A. V. & Salzberg, S. L. The genome polishing tool polca makes fast and accurate corrections in genome assemblies. PLoS Computational Biology 16, e1007981 (2020).

[27] Wick, R. R. & Holt, K. E. Polypolish: short-read polishing of long-read bacterial genome assemblies. PLoS Computational Biology 18, e1009802 (2022).

[28] Vaser, R., Sović, I., Nagarajan, N. & Šikić, M. Fast and accurate de novo genome assembly from long uncorrected reads. Genome Research 27, 737–746 (2017).

[29] Morisse, P., Marchet, C., Limasset, A., Lecroq, T. & Lefebvre, A. Scalable long read self-correction and assembly polishing with multiple sequence alignment. Scientific Reports 11, 761 (2021).

[30] Hu, J. et al. Nextpolish2: a repeat-aware polishing tool for genomes assembled using hifi long reads. Genomics, Proteomics & Bioinformatics 22, qzad009 (2024).

[31] Mastoras, M. et al. Highly accurate assembly polishing with DeepPolisher. Genome Research 35, 1595–1608 (2025).

[32] Li, N. & Stephens, M. Modeling linkage disequilibrium and identifying recombination hotspots using single-nucleotide polymorphism data. Genetics 165, 2213–2233 (2003).

[33] Cleary, J. G. et al. Comparing variant call files for performance benchmarking of next-generation sequencing variant calling pipelines. bioRxiv (2015).

[34] English, A. C., Menon, V. K., Gibbs, R. A., Metcalf, G. A. & Sedlazeck, F. J. Truvari: refined structural variant comparison preserves allelic diversity. Genome Biology 23, 271 (2022).

[35] Garimella, K. V. et al. Population-scale long-read sequencing in the All of Us research program. medRxiv (2025).

[36] Byrska-Bishop, M. et al. High-coverage whole-genome sequencing of the expanded 1000 genomes project cohort including 602 trios. Cell 185, 3426–3440 (2022).

[37] Zook, J. M. et al. Extensive sequencing of seven human genomes to characterize benchmark reference materials. Scientific data 3, 160025 (2016).

[38] Poplin, R. et al. A universal snp and small-indel variant caller using deep neural networks. Nature Biotechnology 36, 983–987 (2018).

[39] Patterson, M. et al. Whatshap: weighted haplotype assembly for future-generation sequencing reads. Journal of Computational Biology 22, 498–509 (2015).

[40] Smolka, M. et al. Detection of mosaic and population-level structural variants with Sniffles2. Nature Biotechnology 42, 1571–1580 (2024).

[41] Rhie, A., Walenz, B. P., Koren, S. & Phillippy, A. M. Merqury: reference-free quality, completeness, and phasing assessment for genome assemblies. Genome Biology 21, 245 (2020).

[42] Schloissnig, S. et al. Structural variation in 1,019 diverse humans based on long-read sequencing. Nature 644, 442–452 (2025).

[43] Išerić, H., Alkan, C., Hach, F. & Numanagić, I. Fast characterization of segmental duplication structure in multiple genome assemblies. Algorithms for Molecular Biology 17, 1–15 (2022).

[44] Smit, A., Hubley, R. & Green, P. RepeatMasker Open-4.0. URL http://www.repeatmasker.org. 2013-2015.

[45] Wagner, J. et al. Curated variation benchmarks for challenging medically relevant autosomal genes. Nature Biotechnology 40, 672–680 (2022).

[46] Taylor-Weiner, A. et al. Scaling computational genomics to millions of individuals with gpus. Genome Biology 20, 228 (2019).

[47] Sahlin, K. Strobealign: flexible seed size enables ultra-fast and accurate read alignment. Genome Biology 23, 260 (2022).

[48] Li, H. Minimap2: pairwise alignment for nucleotide sequences. Bioinformatics 34, 3094–3100 (2018).

[49] Danecek, P. et al. Twelve years of SAMtools and BCFtools. Gigascience 10, giab008 (2021).

