## Supplementary Material for "Scalable and rare-variant aware genome inference across the 1kGP cohort"

June 29, 2026

#### 1 Data

The CHM13-based Minigraph-Cactus data was obtained from:

- GFA: https://s3-us-west-2.amazonaws.com/human-pangenomics/pangenomes/scratch/2025_02_28_minigraph_cactus/hprc-v2.0-mc-chm13/hprc-v2.0-mc-chm13.gfa.gz
- VCF with top-level bubbles: https://s3-us-west-2.amazonaws.com/human-pangenomics/pangenomes/scratch/2025_02_28_minigraph_cactus/hprc-v2.0-mc-chm13/hprc-v2.0-mc-chm13.vcf.gz

Our preprocessing pipeline is availabe as a Snakemake workflow and can be found at: https://github.com/eblerjana/pangenie/tree/master/pipelines/prepare-vcf-from-MC

Links to the PanGenie input VCFs resulting from preprocessing steps described above are provided below. All VCFs are CHM13-based.

- bubble VCF (used as input to PanGenie): https://s3-us-west-2.amazonaws.com/human-pangenomics/pangenomes/scratch/2026_03_30_pangenie/mc_filtered_ids.vcf.gz
- callset VCF (decomposed alleles, used to convert PanGenie genotypes for bubbles to variant genotypes): https://s3-us-west-2.amazonaws.com/human-pangenomics/pangenomes/scratch/2026_03_30_pangenie/mc_filtered_ids_biallelic.vcf.gz

The corresponding reference genome was obtained from: https://s3-us-west-2.amazonaws.com/human-pangenomics/T2T/CHM13/assemblies/analysis_set/chm13v2.0_maskedY_rCRS.fa.gz

Illumina data used for the genotyping experiments described in this paper were downloaded from:

- 3,202 1kGP samples: http://ftp.sra.ebi.ac.uk/vol1/fastq/
- HG002: https://ftp-trace.ncbi.nlm.nih.gov/giab/ftp/data/AshkenazimTrio/HG002_NA24385_son/NIST_Illumina_2x250bps/reads/
- NA21309: https://s3-us-west-2.amazonaws.com/human-pangenomics/index.html?prefix=working/HPRC_PLUS/NA21309/raw_data/Illumina/child/
- HG01123, HG02486, HG02559: https://s3-us-west-2.amazonaws.com/human-pangenomics/index.html?prefix=submissions/30E441F3-6820-4BF6-BCF4-E64D56C8D6A4--TRUSEQ/
- HG005: https://s3-us-west-2.amazonaws.com/human-pangenomics/index.html?prefix=submissions/FFC78D9F-296E-41DD-9E50-C4B5806613EE--HPRC_PLUS_GIAB/HG005/raw_data/Illumina/child/brain-genomics/

#### 2 Haplotype sampling algorithm

We assume each bubble in the pangenome graph to be covered by $N$ haplotypes. We provide an example in Figure 1. Each haplotype traverses one allele of a bubble and multiple haplotypes might traverse the same allele. The mapping between haplotypes and alleles is given by $Allele(h_i)$ (see Figure 1). We define a set of unique k-mers for each bubble containing k-mers that occur exactly once per allele. In the example shown in Figure 1, there are seven unique k-mers. The function $Kmers(a_{j,k})$ returns the set of unique k-mers for a given allele $a_{j,k}$ of a bubble. Similarly, the function $PresentKmers(a_{j,k})$ returns the subset of unique k-mers which additionally have been seen in the sequencing data of the individual that is to be genotyped. Here, we consider a k-mer to be present, if its count in the sequencing reads is at least three.

#### 3 Benchmarking Experiments

We performed leave-one-out experiments to evaluate PanGenie v4. We repeatedly removed one sample from the input VCF provided to PanGenie, genotyped this sample based on the remaining haplotypes in the graph as well as Illumina data, and used the genotypes of the left-out sample as ground truth for evaluation. PanGenie computes genotypes for all bubbles in the input graph. We convert multi-allelic bubble genotypes to bi-allelic genotypes for all nested variants. We then computed the weighted genotype concordance (wGC) as well as precision and recall as a metrics to evaluate the results using the metrics described below.

In addition to leave-one-out experiments, we also genotyped samples not present in the graph and used independent SV callsets from the HGSVC3 as a ground truth for evaluation, which mimicks a more realistic scenario than the leave-one-out setting. We used the PAV calls (`https://ftp.1000genomes.ebi.ac.uk/vol1/ftp/data_collections/HGSVC3/release/Variant_Calls/1.0/T2T-CHM13/variants_T2T-CHM13_sv_insdel_alt_HGSVC2024v1.0.vcf.gz`) as well as the Minigraph-Cactus derived calls (`https://ftp.1000genomes.ebi.ac.uk/vol1/ftp/data_collections/HGSVC3/release/Graph_Genomes/1.0/2024_02_23_minigraph_cactus_hgsvc3/hgsvc3-2024-02-23-mc-chm13-vcfbub.a100k.wave.norm.vcf.gz`) for samples HG00171, NA18989, NA20847, HG02106 and HG02554 as ground truth sets for evaluation. We genotyped these five samples using PanGenie v4 (15 haplotypes) as well as KAGE. We evaluated genotyping results using Truvari [1].

##### 3.1 Running PanGenie

We ran PanGenie v4 (v4.2.1) using the data described in Section 1, using the files and commands shown below. For the leave-one-out experiments, we used the file `mc_filtered_ids.vcf` (see Section 1) and removed the sample to be genotyped from it prior to running the commands below. For the remaining experiments, we directly used the file `mc_filtered_ids.vcf` as the "bubble VCF". We first run the indexing step:

```
PanGenie-index -v <bubble VCF> -r chm13v2.0_masked_rCRS.fa -o index -t 24
```

For genotyping, we ran the following command for each sample:

```
PanGenie -f index -i <sample>_reads.fasta -o pangenie_<sample> -t 24 -j 24 -s <sample>
```

and converted bubble genotypes to genotypes for all nested variants using the script available at `https://github.com/eblerjana/pangenie/blob/master/pipelines/run-from-callset/scripts/convert-to-biallelic.py` and the command shown below.

```
cat pangenie_<sample>_genotyping.vcf | python3 convert-to-biallelic.py mc_filtered_ids_biallelic.vcf.gz |
bgzip > <sample>_genotyping_bi.vcf.gz
```

##### 3.2 Running KAGE

###### 3.2.1 Issues with KAGE2

We first tried running KAGE2 (`https://github.com/kage-genotyper/kage`, commit: 28616ef) but were unsuccessful.

We ran KAGE's indexing step on our decomposed panel VCF (`mc_filtered_ids_biallelic.vcf`) but got the following error: `"ValueError: Mismatching dimensions along axis 1: 1, 462"`. When splitting the input VCF by chromosome, indexing ran through, except for chromosomes chr3, chr4, chr7 and chrX, for which we again got the error above. On the remaining chromosomes, indexing and genotyping (with default parameters) ran successfully without the `--glimpse` option. However, when evaluating the results based on independent PAV and MC callsets for HGSVC3 samples (as done in the manuscript), we observed very low precision for SVs (around 0.40 vs. 0.71 for PanGenie), even inside of GIAB easy regions (0.27 vs. 0.94 for PanGenie). With the `--glimpse` option, the genotyping step fails (see Issue https://github.com/kage-genotyper/kage/issues/9).

We additionally experimented with running KAGE from the "bubble VCF" (`mc_filtered_ids.vcf`), but this again produced the error: `"ValueError: Mismatching dimensions along axis 1: 1, 462"`. We have contacted the developers of KAGE, however, the issues could not be solved by the time of submission of our manuscript. It remains unclear if the problems are related to our data or whether they are caused by bugs in the KAGE software.

##### 3.2.2 AoU KAGE pipeline

Due to the issues described above, we decided to run an older version of KAGE, which was developed in [2] to genotype target samples against a PanGenie-compatible panel of 1,074 samples from the AoU long-read cohort. This version of KAGE was forked from commit 9bcbcd0 of https://github.com/kage-genotyper/kage, which follows v0.1.1 but predates v2.0.0, and introduces relatively minor changes to the original codebase to improve scalability and robustness.

At the pipeline level, one additional methodological change was made to improve scalability; namely, instead of taking an alignment-free approach, aligned reads in CRAM format were taken as input and KAGE was run on a per-chromosome basis. This choice trades scalability for accuracy, since 1) selection of kmers from per-chromosome panels may be less optimal than from a whole-genome panel and 2) signal in unmapped reads may also be lost.

As in [2] (and as originally recommended by the KAGE authors), we follow KAGE genotyping with GLIMPSE v1.1.1 imputation, again running on a per-chromosome basis (although sharding more finely is also possible). KAGE code and WDL pipelines for running KAGE and GLIMPSE are available in commit 2dfb80f of https://github.com/broadinstitute/kage-lite-development and in commit ac33008 of https://github.com/broadinstitute/lrma-aou1-panel-creation, respectively.

#### 3.3 Evaluation metrics

##### 3.3.1 Weighted genotype concordance (wGC)

We reuse our definition of wGC that we had introduced in the original PanGenie publication [3]. We define:

$$\text{weighted genotype concordance} = \frac{\text{conc}(0/0) + \text{conc}(0/1) + \text{conc}(1/1)}{3}$$

where *conc*(0/0), *conc*(0/1) and *conc*(1/1) denote the fractions of all 0/0, 0/1 and 1/1 ground truth genotypes correctly genotyped as such by the genotyper, respectively. Included are all graph variants that are genotypable, that is, we exclude variants that are unique to the evaluation sample. These variants are not part of the set of input variants given to the genotypers and thus will not be considered for genotyping (as the tools re-genotype variants and do not detect them) [3].

##### 3.3.2 Precision and recall

In addition to the wGC metric, we also used RTG vcfeval [4] to evaluate small variants (SNPs and indels $< 50$ bp), as well as Truvari [1] to evaluate SVs $\geq 50$ bp. Both tools report precision, recall and F-score metrics. While vcfeval is genotype-aware and requires correct genotypes to define true positive variants, Truvari only considers presence or absence signals, that is, it does not distinguish 0/1 and 1/1 genotypes. It additionally outputs its own genotype concordance metric which measures the fraction of matching genotypes within the set of true positive variants. We compute *adjusted* versions of precision and F-scores by excluding variants from the ground truth set which are unique to the sample to be genotyped and thus not genotypable by any re-genotyper.

#### 3.4 Evaluation regions

Besides evaluating genotyping results across the whole genome, we also stratified by different regions. We used the following regions defined by GIAB (https://www.nist.gov/programs-projects/genome-bottle, v3.3):

- **Easy regions** obtained from: https://ftp-trace.ncbi.nlm.nih.gov/ReferenceSamples/giab/release/genome-stratifications/v3.3/CHM13@all/Union/CHM13_notinalldifficultregions.bed.gz

- **SegDup regions** obtained from: https://ftp-trace.ncbi.nlm.nih.gov/ReferenceSamples/giab/release/genome-stratifications/v3.3/CHM13@all/Union/CHM13_alllowmapandsegdupregions.bed.gz

- **Repeat regions** obtained from: https://ftp-trace.ncbi.nlm.nih.gov/ReferenceSamples/giab/release/genome-stratifications/v3.3/CHM13@all/LowComplexity/CHM13_AllTandemRepeatsandHomopolymers_slop5.bed.gz

All regions are relative to the CHM13 reference genome.

#### 3.5 Benchmarking different sampling sizes

Our haplotype sampling algorithm generates a smaller subset of haplotype paths from the full set of haplotypes present in the pangenome graph provided as an input to PanGenie. Our sampling algorithm is iterative and has to be run $R$-times ($R << N$) in order to produce a set of $R$ haplotype sequences. We have experimented with different parameters for $R$ in order to analyze runtime and memory consumption of PanGenie, as well as the resulting genotyping accuracy. For this purpose, we performed a leave-one-out experiment based on the HPRC v2.0 data, with varying numbers of sampled haplotypes. Results are shown in Figure 2 of the manuscript as well as in Supplementary Figures 2 and 3. PanGenie v4 tends to outperform PanGenie v4 (chunking), and increasing the number of sampled haplotypes often increases accuracy. At the same time, runtime and memory usages are much lower for PanGenie v4 (Figure 2 and Supplementary Figure 4).

#### 3.6 Comparison to the HGSVC3 graph

We additionally performed the same leave-one-out experiment but with using the HGSVC3 pangenome reference containing 216 haplotypes (obtained from: https://ftp.1000genomes.ebi.ac.uk/vol1/ftp/data_collections/HGSVC3/release/Genotyping_1kGP/PanGenie-genotypes/1.0/). We compared genotyping performances for PanGenie v4 (with 15 sampled haplotypes) on both data sets (HPRC2 and HGSVC3). Results are shown in Supplementary Figure 5. Results show better genotyping performance on the new HPRC v2.0 graph. The only exception is sample HG01891 which performs better with the HGSVC3 graph. This, however, can be explained by the fact that the HGSVC3 graph contains one of its parent samples, HG01890 while the HPRC v2.0 graph does not.

##### 3.6.1 Comparison to other genotyping methods

We genotyped five independent samples (HG00171, NA18989, NA20847, HG02106, HG02554), which are not in the HPRC v2.0 graph, using PanGenie v4 and KAGE and evaluated the resulting geno- types based on independent ground truth SV calls produced by the HGSVC for these samples [5]. These ground truth sets included a PAV callset generated directly from haplotype-resolved assemblies of these samples as well as a Minigraph-Cactus callset generated from a pangenome graph of the HGSVC assemblies [5]. We evaluated SNPs and indels with vcfeval [4] and SVs with Truvari [1]. We stratified results by different regions 3.4.

Results for SNPs + indels evaluated based on both ground truth sets are shown in Supplementary Figures 6 and 7. For SVs, results are shown in Supplementary Figures 8 and 9.

#### 3.7 Runtimes

All PanGenie experiments were run on a HPC-cluster predominately consisting of Intel E5-2697v2 (2 × 12 cores and 128 GB of RAM) and Intel Xeon Gold 6136 (2 × 12 cores and 192 GB of RAM) nodes. We provide runtimes of PanGenie v4 in Supplementary Table 1.

All KAGE+GLIMPSE experiments were run on the Terra cloud-computing platform. For the KAGE pipeline, average walltime (cloud cost) per sample was 63 minutes ($0.31); besides KAGE kmer counting and genotyping, this compute time/cost also encompasses CRAM decompression, virtual-machine (VM) preemptions, and Terra job queuing. For each sample, the cost-dominating step in this pipeline was sharded across 4 VMs, each with 6 vCPUs and 6GB and covering roughly equal portions of the genome. In contrast, for the GLIMPSE pipeline, all five samples were run in a single batch job with a walltime (cloud cost) of 136 minutes ($2.77). The dominating step in the GLIMPSE pipeline was the running of the GLIMPSE `phase` tool, which was sharded across per-chromosome VMs, each with 6 vCPUs and memory scaling linearly with chromosome size from 20-100GB. These VMs were slightly overprovisioned, as they were only roughly adjusted from the configurations used to run larger batches of AoU samples in [2] and were not cost optimized for the batch under consideration.

| sample | PanGenie v4 | |
|---|---|---|
| | CPU time (s) | walltime (hh:mm:ss, 24 cores) |
| HG00171 | 34,702 | 0:53:28 |
| HG02106 | 38,293 | 0:56:37 |
| HG02554 | 40,647 | 0:59:54 |
| NA18989 | 46,376 | 1:04:08 |
| NA20847 | 32,088 | 0:49:47 |

Table 1: **Runtimes of PanGenie.**

### 4 Genotyping the 1kGP cohort

We used the VCF file derived from the HPRC v2.0 graph to genotype all 3,202 samples that are part of the 1kGP cohort as well as 9 additional HPRC samples that are not overlapping the 1kGP set (HG01123, HG02486, HG02559, NA21309, HG002, HG02109, HG03471, HG06807, HG005) with PanGenie v4.2.1 based on high coverage Illumina data [6, 7, 8]. We ran PanGenie the same way as described in Section 3.1. More specifically, we used the command below for indexing (run once) using the data described in Section 1.

```
PanGenie-index -v mc_filtered_ids.vcf -r chm13v2.0_masked_rCRS.fa -o index -t 24
```

For genotyping, we ran the following command for each sample:

```
PanGenie -f index -i <sample>_reads.fasta -o pangenie_<sample> -t 24 -j 24 -s <sample>
```

and converted bubble genotypes to genotypes for all nested variants using the script available at https://github.com/eblerjana/pangenie/blob/master/pipelines/run-from-callset/scripts/convert-to-biallelic.py and the command shown below.

```
cat pangenie_<sample>_genotyping.vcf | python3 convert-to-biallelic.py mc_filtered_ids_biallelic.vcf.gz |
bgzip > <sample>_genotyping_bi.vcf.gz
```

We ran our previously developed filtering pipeline [9, 8, 5] based on a regression model to filter the genotypes. Compared to our previous work [5], we used slightly modified thresholds for our filters (listed below), but otherwise used the same regression model that we have described previously [5].

- **ac0_fail**: a variant allele was genotyped with a allele frequency of 0.0 across all samples
- **mendel_fail**: the mendelian consistency across trios is less than 85 % for a variant allele. Here, we use a strict definition of mendelian consistency which excludes all trios with only 0/0, only 0/1 and only 1/1 genotypes.
- **gq_fail**: less than 20 high quality genotypes were reported for this variant allele
- **self_fail**: genotyping accuracy of a variant allele across the panel samples is less than 90%
- **nonref_fail**: not a single non-0/0 genotype was genotyped correctly across all panel samples

Our filtered set of variants contained 39,366,061 of the 46,996,181 SNPs + indels (83.8 %), 117,555 of 127,964 SV deletions (92%), 597,814 of 643,535 SV insertions (93%) and 117,359 of 129,530 other SV alleles (91 %). We compared allele frequencies of our genotyped variants (before and after filtering) with the allele frequency of the variants in the assembly (i.e. the input HPRC panel). Before filtering, we observed correlations of 0.993, 0.992, 0.991, 0.987 and 0.965 for SNPs, indels, SV deletions, SV insertions and other SV alleles, respectively. After filtering, we observed correlations of 0.998, 0.998, 0.997, 0.994 and 0.982 (Supplementary Figure 10 a, Figure 3a). Heterozygosities and allele frequencies of the genotyped variants follow the Hardy-Weinberg equilibrium (Supplementary Figure 10 b, Figure 3b).

We compared our filtered genotyped variants for the 3,202 1kGP samples to two other SV sets for the same cohort: our previously generated PanGenie genotypes produced for the HGSVC3 project, produced by genotyping variants detected across 216 assembled haplotypes across the 1kGP cohort (https://ftp.1000genomes.ebi.ac.uk/vol1/ftp/data_collections/HGSVC3/release/Genotyping_1kGP/PanGenie-genotypes/1.0/pangenie_chm13_all_decomposed_lenient.vcf.gz) as well as an Illumina based SV discovery callset generated by alignment-based SV detection tools across all 3,202 samples (https://ftp.1000genomes.ebi.ac.uk/vol1/ftp/data_collections/1000G_2504_high_coverage/working/20210124.SV_Illumina_Integration/1KGP_3202.gatksv_svtools_novelins.freeze_V3.wAF.vcf.gz). Since the latter calls are relative to GRCh38 and since SV comparison is typically difficult due to representation differences, we compared the callsets based on the number of SVs detected by each sample (Figure 3c). Prior to analysis, we merged similar alleles in both our PanGenie callsets using Truvari's collapse command [1] with parameters: `truvari collapse -r 500 -p 0.95 -P 0.95 -s 50 -S 100000`. Comparison results indicate that we can now capture more SVs per sample with our latest PanGenie set based on the HPRC v2.0 graph. Improvements over the HGSVC3 set are likely a result of better assemblies produced by the HPRC, a larger set of assembled haplotypes in the graph (462 instead of 216) as well as improved genotyping performance with PanGenie v4 (the HGSVC3 genotypes were produced with PanGenie using the chunking strategy).

We proceeded with phasing the unfiltered set of genotypes with SHAPEIT5 (v5.1.1) [10]. We provided the following input data to SHAPEIT: we started from our biallelic VCF (`mc_filtered_ids_biallelic.vcf.gz`, see Section 1) and removed all variants which contained any missing genotype information across the 462 graph haplotypes (since SHAPEIT5 cannot handle them). The resulting VCF was used as a reference panel for phasing (`--reference` parameter). Our unfiltered genotypes across the 1kGP samples were provided to `--input`. We further provided pedigree information (`--pedigree`) and genetic maps obtained from https://github.com/JosephLalli/phasing_T2T/tree/main/resources/recombination_maps/t2t_native_scaled_maps/ (`--map`). SHAPEIT5 was run separately on each chromosome. For chromosome X, we additionally provided a list with all male samples to `--haploid`. Otherwise, we used default parameters.

After phasing, we constructed consensus haplotypes for all samples. For each sample, we generated two FASTA sequences by implanting phased variant alleles into the CHM13 reference genome using the `bcftools consensus` command [11].

### 5 Haplotype polishing

#### 5.1 Polishing with different read data

We have tested our polishing pipeline using different sources of long read data: ONT data produced by Schloissnig et al. [12] ("ONT-S"), ONT data produced by Gustafson et al. [13] ("ONT-G") and HiFi data produced by the HGSVC [5]. Since there was no common sample for which all three data types where available, we ran the polishing workflow on sample HG02282 using both ONT datasets, and on sample HG00096 using the ONT-S and HiFi data. We additionally ran a re-polishing experiment, in which we polished twice, by re-applying the same pipeline to polished haplotypes resulting from the first round. In addition to testing different long read datasets, we also evaluated the impact of the different variant types on the polishing results. For this purpose, we ran three different versions of our polishing pipeline which correct (a) only SNPs, (b) SNPs and indels and (c) SNPs, indels and SVs. Results are shown in Supplementary Figure 11 and underlying numbers of variants incorporated into the haplotypes in each round in Supplementary Table 2. Polishing increases k-mer based as well as variant-based QVs, especially when indels and SVs are incorporated. We furthermore observed better polishing results using HiFi data compared to ONT, which is expected due to the higher accuracy of HiFi reads. Furthermore, running another round of polishing improves the results, which is likely because of more accurate read alignments resulting from using a more accurate consensus sequence as a reference. Since using the combined sets of SNPs, indels and SVs for polishing produced the best results, we used this version of the pipeline for polishing the 1kGP samples. We used ONT-S reads for all 967 samples overlapping with the 1kGP cohort for polishing.

#### 5.2 Evaluating consensus haplotypes

For evaluation, we selected 10 samples (HG02595, HG03091, NA19717, HG01188, HG01595, HG00632, HG00324, NA12890, HG03697, HG03760) which were neither part of the HGSVC3 nor the HPRC v2.0 individuals. This enables a fair comparison of our consensus haplotypes to the ones we had previously generated for the HGSVC3 graph using a similar pipeline. The major difference between our unpolished HPRC consensus haplotypes and those generated for the HGSVC3 is that the latter additionally includes rare SNP and indels called from 1kGP Illumina data [5]. We additionally compared our haplotypes to consensus haplotypes we generated from phased SNP, indel and SV calls generated with traditional alignment-based methods from Illumina data of the 1kGP cohort samples [6]. Since these calls were relative to GRCh38, we used the GRCh38 reference genome when implanting variants. We evaluated the consensus haplotypes by computing k-mer based QV estimates based on Illumina reads

| | HG00096 hap1 | | HG02282 hap1 | |
|---|---|---|---|---|
| | SNPs+indels | SVs | SNPs+indels | SVs |
| ONT-S | 605,778 | 414 | - | - |
| ONT-S repolished | 187,289 | 334 | - | - |
| HiFi | 526,711 | 1,281 | - | - |
| HiFi repolished | 160,288 | 817 | - | - |
| ONT-G | - | - | 639,822 | 1,080 |
| ONT-G repolished | - | - | 185,058 | 615 |

Table 2: **Variants corrected during polishing** Shown are the number of small variants (SNPs and indels) and the number of structural variants that are incorporated into the haplotypes during polishing. Results are shown for one haplotype of two samples (HG00096 and HG02282) polished with different read data. Two rounds of polishing were performed, "repolished" refers to the results after the second round of polishing.

(see below). To ensure a fair comparison between CHM13-based and GRCh38-based haplotypes, we additionally restricted evaluation to syntenic regions, including all regions shared between T2T-CHM13 and GRCh38.

We furthermore selected 8 HGSVC3 samples (HG00096, HG00171, HG01596, HG02554, HG02953, HG03009, NA18989 and NA20847) which do not overlap with our HPRC v2.0 graph and used the HGSVC3 assemblies for these samples as ground truth for evaluation. Since k-mer based QV estimates cannot fully capture structural correctness of our haplotypes, we computed variant-based haplotypes (see below) based on the HGSVC3 assemblies. This also provides local QV estimates for windows along the haplotypes allowing us to analyze accuracy of certain regions of the genome. We show histograms of window-wise QVs for HG03009 computed for 1kGP, HGSVC3 and (un)polished HPRC consensus haplotypes in Figure 12. We furthermore show the distributions of variant-based QVs across all samples in Figure 13. We observed lowest QVs for the 1kGP based haplotypes, and highest QVs for the polished HPRC haplotypes. Median QVs were: 26.4, 33.8, 35.0 and 35.9 for the 1kGP, HGSVC3, unpolished HPRC and polished HPRC consensus haplotypes.

#### 5.3 Evaluation metrics

##### 5.3.1 Variant based QVs

We have previously presented a pipeline to compute variant-based QV estimates [5]. Briefly, our pipeline compares the two consensus haplotype sequences computed for a sample to haplotype-resolved assemblies available for the same individual. We use one of the consensus haplotypes as a reference, the other one as a query, together with the two ground truth haplotype-resolved assemblies of the same sample [5]. We then call variants (SNPs, indels and SVs) using PAV [9] and determine conflicting calls between consensus haplotypes and assemblies [5]. QVs are then computed based on the induced basepair changes of these variants within windows of 1Mbp along the reference haplotype using: $-10 \cdot log_{10}(\text{basepair\_changes}/(2 \cdot \text{window\_size}))$ [5]. We additionally run BISER (v1.4) [14] and RepeatMasker (`http://www.repeatmasker.org/`) on the consensus haplotype sequence and intersect resulting annotation intervals with our 1 Mbp windows.

##### 5.3.2 K-mer based QVs

We computed k-mer based QV estimates using meryl (`https://github.com/marbl/meryl`) analogously to how they are computed by Merqury [15]. Briefly, the idea is to analyze the overlap of k-mers present in the haplotype sequence and the k-mers present in the Illumina reads of the same individual. The higher this overlap, the larger the resulting QV value. We used k-mer size 21 for estimating QVs.

#### 5.4 Runtimes

We show the total single-core runtimes (in CPU seconds) for polishing each haplotype based on different read datasets, starting from the unpolished consensus sequence of each haplotype in Supplementary Table 3. For all these runs, the majority of the runtime is spend for read alignment of Illumina and ONT data and small variant calling with DeepVariant. For sample HG00096 (hap1) using ONT-S data, 42 % of the total runtime is spent aligning reads and another 51 % are spent for small variant calling with DeepVariant. Numbers look similar for other samples and data sets (Supplementary Table 4). In order to decrease the walltime of running our polishing pipeline, we parallelized all steps subsequent of read alignment by chromosome.

| | HG00096 hap1 | HG00096 hap2 | HG02282 hap1 | HG02282 hap2 |
|---|---|---|---|---|
| ONT-S | 572,110 sec | 572,556 sec | - | - |
| HiFi | 500,519 sec | 507,237 sec | - | - |
| ONT-G | - | - | 760,577 sec | 764,448 sec |

Table 3: **Runtime of polishing workflow.** Shown are single-core runtimes (in CPU seconds) for polishing haplotypes of samples HG00096 and HG02282 based on ONT and HiFi data sets.

| sample | time alignment | time DeepVariant | time remaining steps |
|---|---|---|---|
| HG00096 hap1 ONT-S | 239,042 sec | 293,440 sec | 39,628 sec |
| HG00096 hap1 HiFi | 167,554 sec | 291,286 sec | 41,679 sec |
| HG02282 hap1 ONT-G | 370,566 sec | 340,296 sec | 49,715 sec |

Table 4: **Runtimes for different polishing steps.** Shown are single-core runtimes (in CPU seconds) for polishing haplotypes of samples HG00096 and HG02282 based on ONT and HiFi data sets. Runtimes are split by alignment, small variant calling with DeepVariant and the remaining steps.

### 6 Pangenome cis-eQTL mapping in Geuvadis LCL samples

#### 6.1 Cohort and expression data

TMM-normalized RNA-seq expression matrices were obtained from Ebert et al. 2021 [9] for Geuvadis LCL samples [16]. A harmonized subset of 397 individuals with matched short-read WGS was retained after quality control. Genes with expression $> 1$ TPM in $\geq 20\%$ of samples and maximum expression $\geq 1$ TPM were retained; after intersecting with available coordinates, 14,164 genes were carried forward to mapping. Sixty expression PCs were included as fixed-effect covariates.

#### 6.2 Pangenome genotyping

Each of the 397 samples was genotyped independently with PanGenie v4.2.1 against two pangenome variant panels: HPRC v2.0 MC GRCh38 (231 individuals, 462 haplotypes) and HGSVC2 PAV Freeze4 GRCh38 (32 individuals, 64 haplotypes). Two individuals are shared between panels; 20 and 3 Geuvadis samples appear in the HPRC and HGSVC2 panels, respectively. Paired-end reads were streamed to PanGenie via standard input (`-j 24 -t 24`) using a pre-built chromosome-level index.

#### 6.3 Cohort VCF construction and genotype filtering

Per-sample multiallelic VCFs were converted to biallelic representation with `convert-to-biallelic.py` from the PanGenie run-from-callset pipeline. Per-chromosome cohort BCFs were produced with `bcftools merge` and concatenated with `bcftools concat`. Because HPRC pangenome graph-path identifiers exceed PLINK2's 16,384-character limit, variant IDs were replaced with sequential identifiers (`HPRC_1`, `HPRC_2`, ...) with a corresponding map file retained. Cohort BCFs were converted to PLINK2 pgen format (`--bcf --make-pgen --split-par hg38`) and filtered to MAF $\geq 0.01$, yielding 18.4 M variants for HPRC and 11.8 M for HGSVC2.

#### 6.4 cis-eQTL mapping

cis-eQTL mapping was performed with tensorQTL [17] using the permutation pass (`cis.map_cis`) with a $\pm 1$ Mb cis-window and MAF threshold $\geq 0.01$. Per-gene beta-approximated permutation $p$-values were corrected for multiple testing using Storey–Tibshirani $q$-values ($\lambda = 0.85$); genes with $q \leq 0.05$ were declared eGenes (FDR 5%).

##### 6.5 Cross-panel comparison and variant annotation

eGenes were compared between panels by phenotype identifier. Variant type (SNV, INS, DEL, COMPLEX) was extracted from the third hyphen-delimited field of PanGenie variant IDs; allele size was inferred from the numeric suffix (HPRC) or from REF/ALT allele length difference (HGSVC2). Variants with allele size $\geq 50$ bp were classified as large SVs. For each shared eGene, the lead variant type from each panel was compared, and the panel showing the larger $|\hat{\beta}|$ was recorded. Among shared eGenes where HPRC assigned an indel/SV lead and HGSVC2 an SNV lead, the fraction with the larger HPRC $|\hat{\beta}|$ was compared to that in an SNV-concordant comparison set by two-proportion $\chi^2$ test (60.4% vs. 40.0%; $\chi^2 = 273$, $p < 10^{-60}$).

### References


[1] Adam C English, Vipin K Menon, Richard A Gibbs, Ginger A Metcalf, and Fritz J Sedlazeck. Truvari: refined structural variant comparison preserves allelic diversity. *Genome Biology*, 23(1):271, 2022.

[2] Kiran V Garimella, Qiuhui Li, Julie Wertz, Samuel K Lee, Fabio Cunial, Yongqing Huang, Yulia Mostovoy, Ryan Lorig-Roach, Adam English, Hang Su, et al. Population-scale long-read sequencing in the all of us research program. *medRxiv*, 2025.

[3] Jana Ebler, Peter Ebert, Wayne E Clarke, Tobias Rausch, Peter A Audano, Torsten Houwaart, Yafei Mao, Jan O Korbel, Evan E Eichler, Michael C Zody, et al. Pangenome-based genome inference allows efficient and accurate genotyping across a wide spectrum of variant classes. *Nature genetics*, 54(4):518–525, 2022.

[4] John G Cleary, Ross Braithwaite, Kurt Gaastra, Brian S Hilbush, Stuart Inglis, Sean A Irvine, Alan Jackson, Richard Littin, Mehul Rathod, David Ware, et al. Comparing variant call files for performance benchmarking of next-generation sequencing variant calling pipelines. *BioRxiv*, page 023754, 2015.

[5] Glennis A Logsdon, Peter Ebert, Peter A Audano, Mark Loftus, David Porubsky, Jana Ebler, Feyza Yilmaz, Pille Hallast, Timofey Prodanov, DongAhn Yoo, et al. Complex genetic variation in nearly complete human genomes. *Nature*, pages 1–12, 2025.

[6] Marta Byrska-Bishop, Uday S Evani, Xuefang Zhao, Anna O Basile, Haley J Abel, Allison A Regier, André Corvelo, Wayne E Clarke, Rajeeva Musunuri, Kshithija Nagulapalli, et al. High-coverage whole-genome sequencing of the expanded 1000 genomes project cohort including 602 trios. *Cell*, 185(18):3426–3440, 2022.

[7] Justin M Zook, David Catoe, Jennifer McDaniel, Lindsay Vang, Noah Spies, Arend Sidow, Ziming Weng, Yuling Liu, Christopher E Mason, Noah Alexander, et al. Extensive sequencing of seven human genomes to characterize benchmark reference materials. *Scientific data*, 3(1):160025, 2016.

[8] Wen-Wei Liao, Mobin Asri, Jana Ebler, Daniel Doerr, Marina Haukness, Glenn Hickey, Shuangjia Lu, Julian K Lucas, Jean Monlong, Haley J Abel, et al. A draft human pangenome reference. *Nature*, 617(7960):312–324, 2023.

[9] Peter Ebert, Peter A Audano, Qihui Zhu, Bernardo Rodriguez-Martin, David Porubsky, Marc Jan Bonder, Arvis Sulovari, Jana Ebler, Weichen Zhou, Rebecca Serra Mari, et al. Haplotype-resolved diverse human genomes and integrated analysis of structural variation. *Science*, 372(6537):eabf7117, 2021.

[10] Robin J Hofmeister, Diogo M Ribeiro, Simone Rubinacci, and Olivier Delaneau. Accurate rare variant phasing of whole-genome and whole-exome sequencing data in the uk biobank. *Nature genetics*, 55(7):1243–1249, 2023.

[11] Petr Danecek, James K Bonfield, Jennifer Liddle, John Marshall, Valeriu Ohan, Martin O Pollard, Andrew Whitwham, Thomas Keane, Shane A McCarthy, Robert M Davies, et al. Twelve years of samtools and bcftools. *Gigascience*, 10(2):giab008, 2021.

[12] Siegfried Schloissnig, Samarendra Pani, Jana Ebler, Carsten Hain, Vasiliki Tsapalou, Arda Söylev, Patrick Hüther, Hufsah Ashraf, Timofey Prodanov, Mila Asparuhova, et al. Structural variation in 1,019 diverse humans based on long-read sequencing. *Nature*, pages 1–11, 2025.

[13] Jonas A Gustafson, Sophia B Gibson, Nikhita Damaraju, Miranda PG Zalusky, Kendra Hoekzema, David Twesigomwe, Lei Yang, Anthony A Snead, Phillip A Richmond, Wouter De Coster, et al. High-coverage nanopore sequencing of samples from the 1000 genomes project to build a comprehensive catalog of human genetic variation. *Genome research*, 34(11):2061–2073, 2024.

[14] Hamza Išerić, Can Alkan, Faraz Hach, and Ibrahim Numanagić. Fast characterization of segmental duplication structure in multiple genome assemblies. *Algorithms for Molecular Biology*, 17(1):1–15, 2022.

[15] Arang Rhie, Brian P Walenz, Sergey Koren, and Adam M Phillippy. Merqury: reference-free quality, completeness, and phasing assessment for genome assemblies. *Genome biology*, 21(1):245, 2020.

[16] Tuuli Lappalainen, Michael Sammeth, Marc R Friedländer, Peter AC 't Hoen, Jean Monlong, Manuel A Rivas, Mar Gonzalez-Porta, Natalja Kurbatova, Thasso Griebel, Pedro G Ferreira, et al. Transcriptome and genome sequencing uncovers functional variation in humans. *Nature*, 501(7468):506–511, 2013.

[17] Amaro Taylor-Weiner, François Aguet, Nicholas J Haradhvala, Sager Gosai, Shankara Anand, Jaegil Kim, Kristin Ardlie, Eliezer M Van Allen, and Gad Getz. Scaling computational genomics to millions of individuals with gpus. *Genome Biology*, 20(1):228, 2019.


### Supplementary Figures

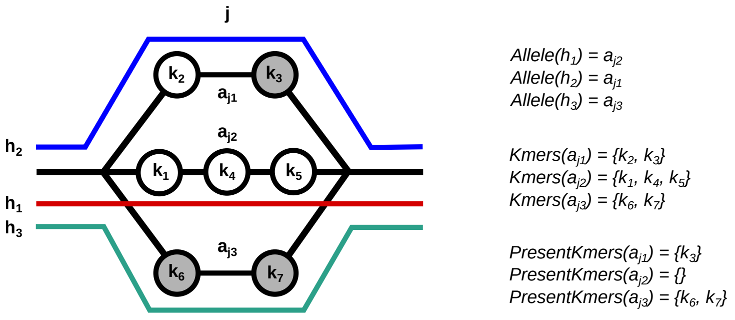


Figure 1: **Definitions** Example of a bubble $j$ with three different alleles covered by three haplotypes. Circles represent unique k-mers, the ones colored in grey represent the k-mers which are present in the sequencing reads of the sample to be genotyped.

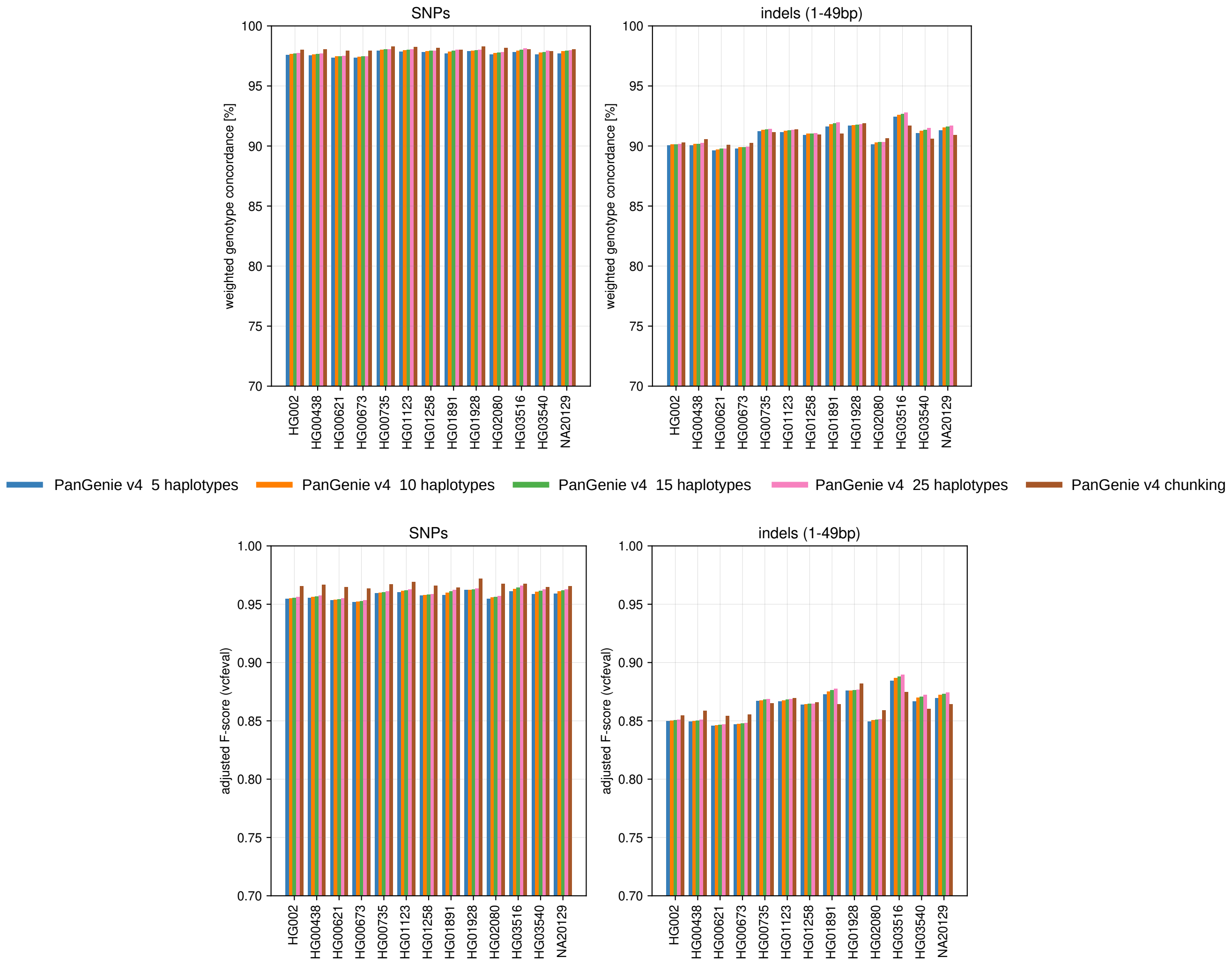


Figure 2: **Number of sampled haplotypes - SNPs + indels** F-scores for genotypes computed with different numbers of haplotypes sampled in the haplotype sampling step implemented in PanGenie v4. Additionally, we ran PanGenie with the previously implemented chunking strategy instead of haplotype sampling.

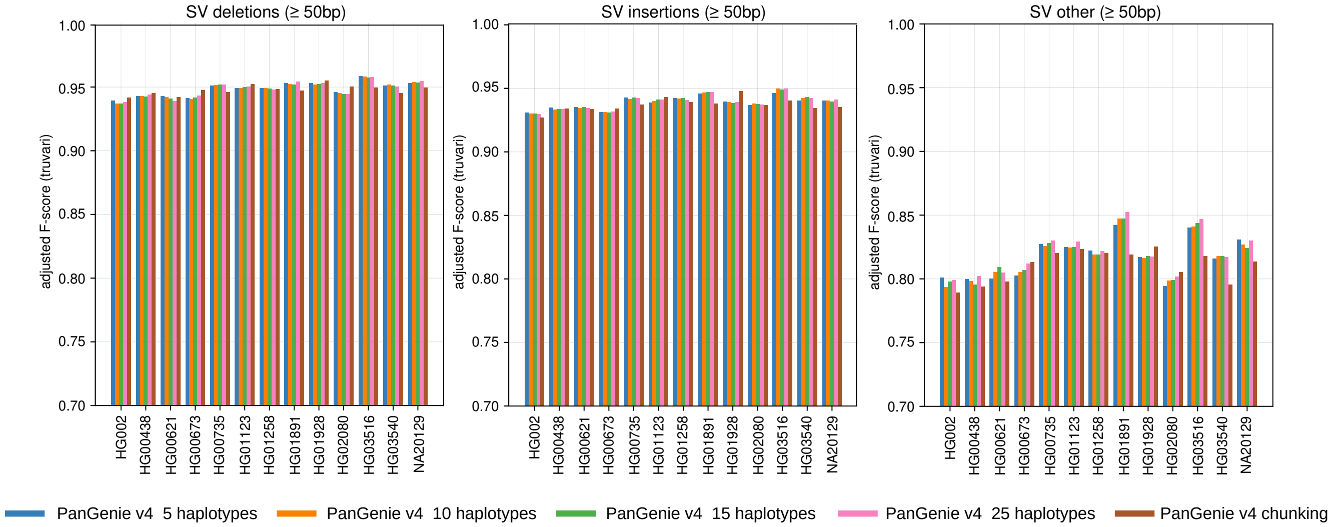


Figure 3: **Number of sampled haplotypes - SVs** F-scores for genotypes computed with different numbers of haplotypes sampled in the haplotype sampling step implemented in PanGenie v4. Additionally, we ran PanGenie with the previously implemented chunking strategy instead of haplotype sampling.

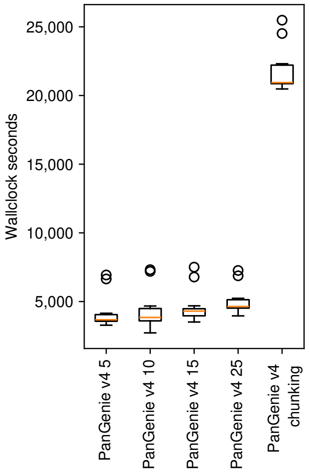


Figure 4: **Runtimes** Wallclock times of PanGenie v4 with chunking and PanGenie v4 with different number of sampled haplotypes (5-25) across 13 samples with 24 cores.

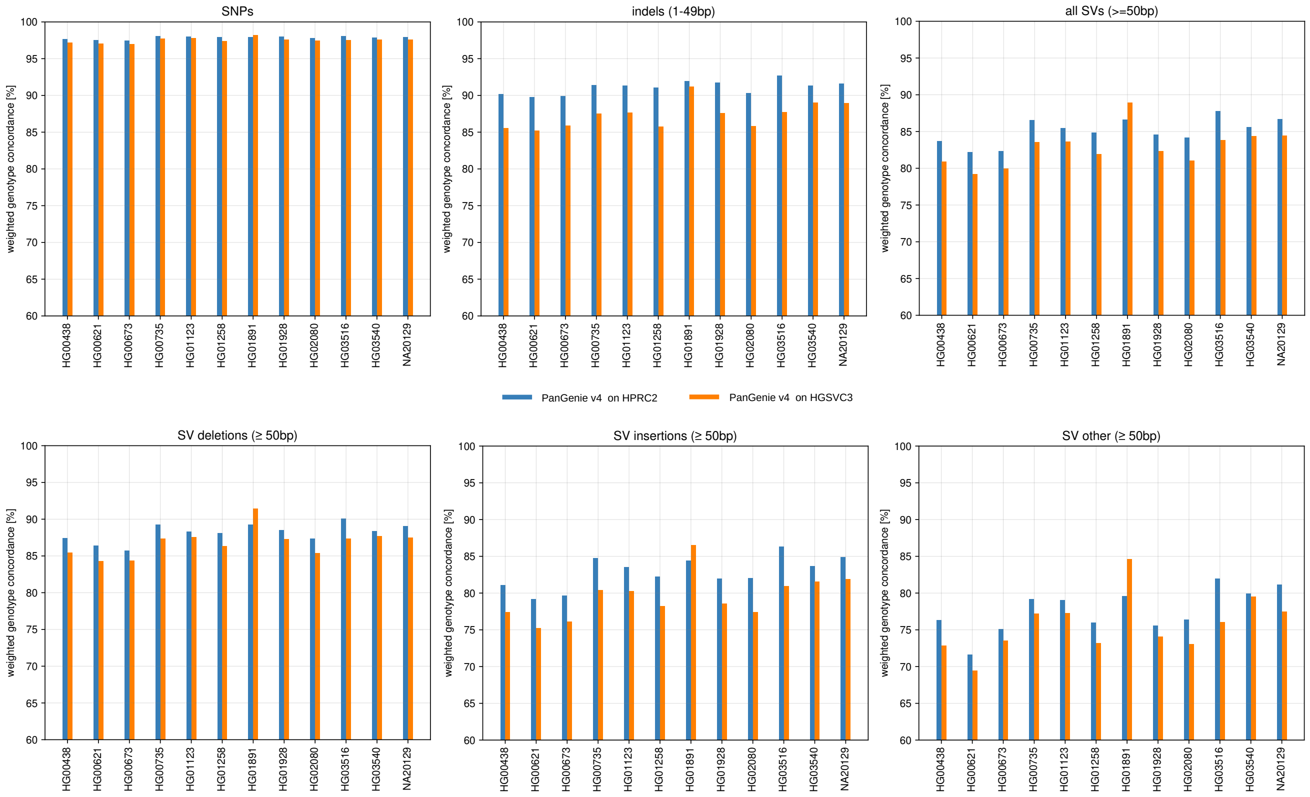


Figure 5: **Comparison to HGSVC3** Leave-one-out experiments were performed, in which one sample was repeatedly taken out of the graph, genotyped based on the remaining samples with PanGenie (v4, 15 haplotypes), and the left-out genotypes were used as ground truth for evaluation. We performed this experiment for 12 samples and ran it twice: once using the HPRC v2.0 graph containing 462 haplotypes and once using the HGSVC3 graph containing 216 haplotypes. Note that for sample HG01891, the HGSVC3 graph contains a parent sample (HG01890), while the HPRC graph does not. This is a likely explanation for the superior performance of this sample on the HGSVC3 panel.

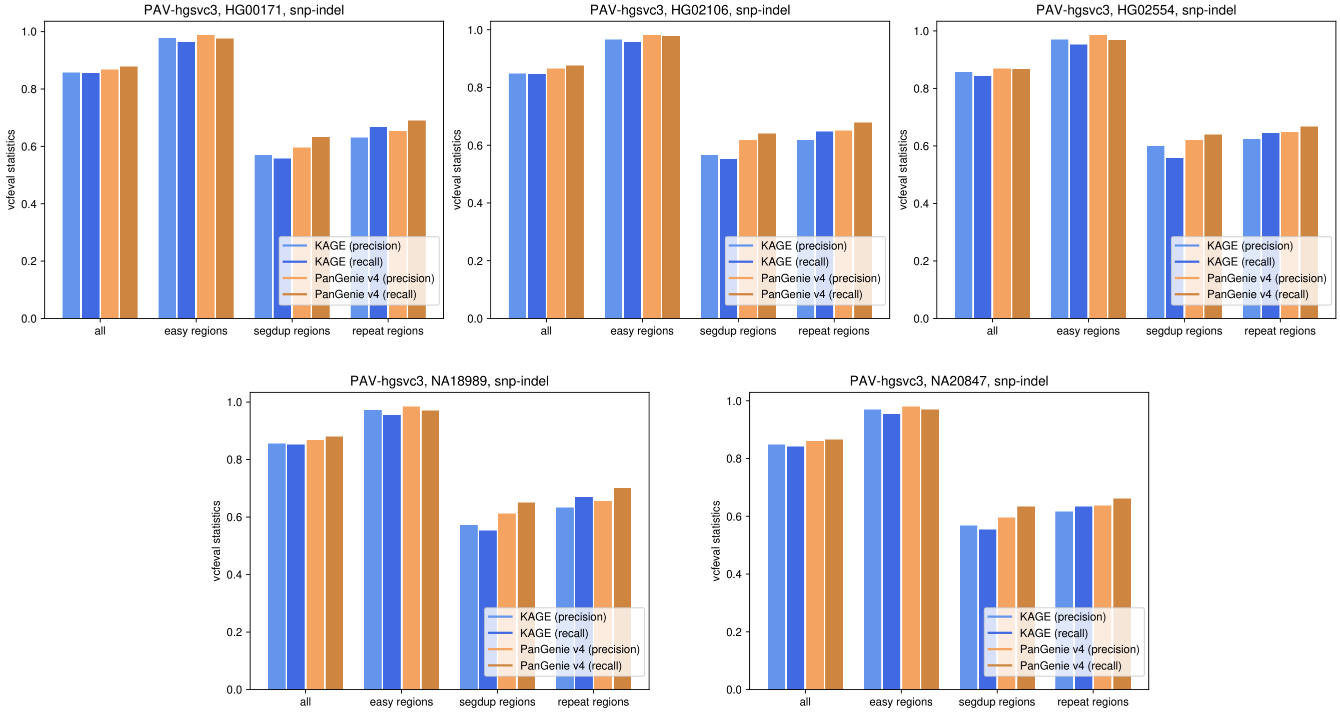


Figure 6: **Evaluation of SNPs and indels with PAV ground truth** Precision and recall statistics for PanGenie and KAGE computed for five samples based on the HGSVC3 PAV ground truth SNP and indel calls. Results are stratified by GIAB regions.

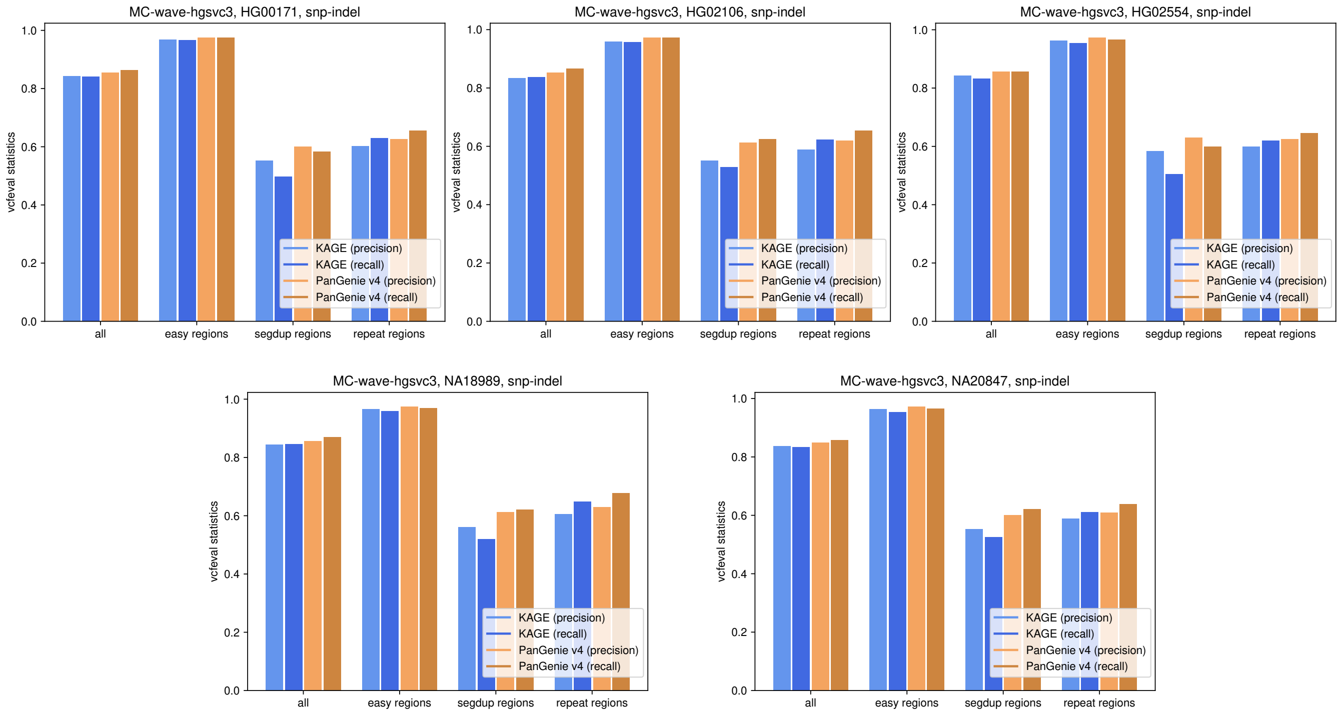


Figure 7: **Evaluation of SNPs and indels with MC ground truth** Precision and recall statistics for PanGenie and KAGE computed for five samples based on the HGSVC3 Minigraph-Cactus ground truth SNP and indel calls. Results are stratified by GIAB regions.

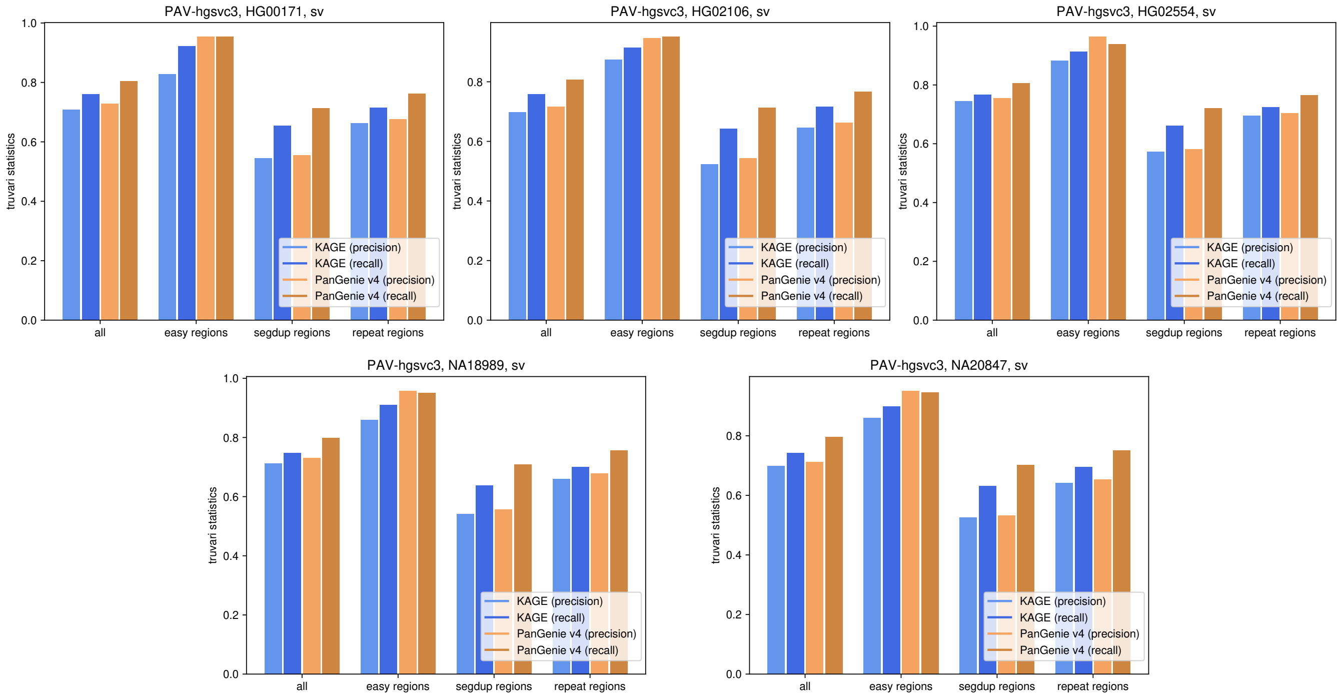


Figure 8: **Evaluation of SVs with PAV ground truth** Precision and recall statistics for PanGenie and KAGE computed for five samples based on the HGSVC3 PAV ground truth SV calls. Results are stratified by GIAB regions.

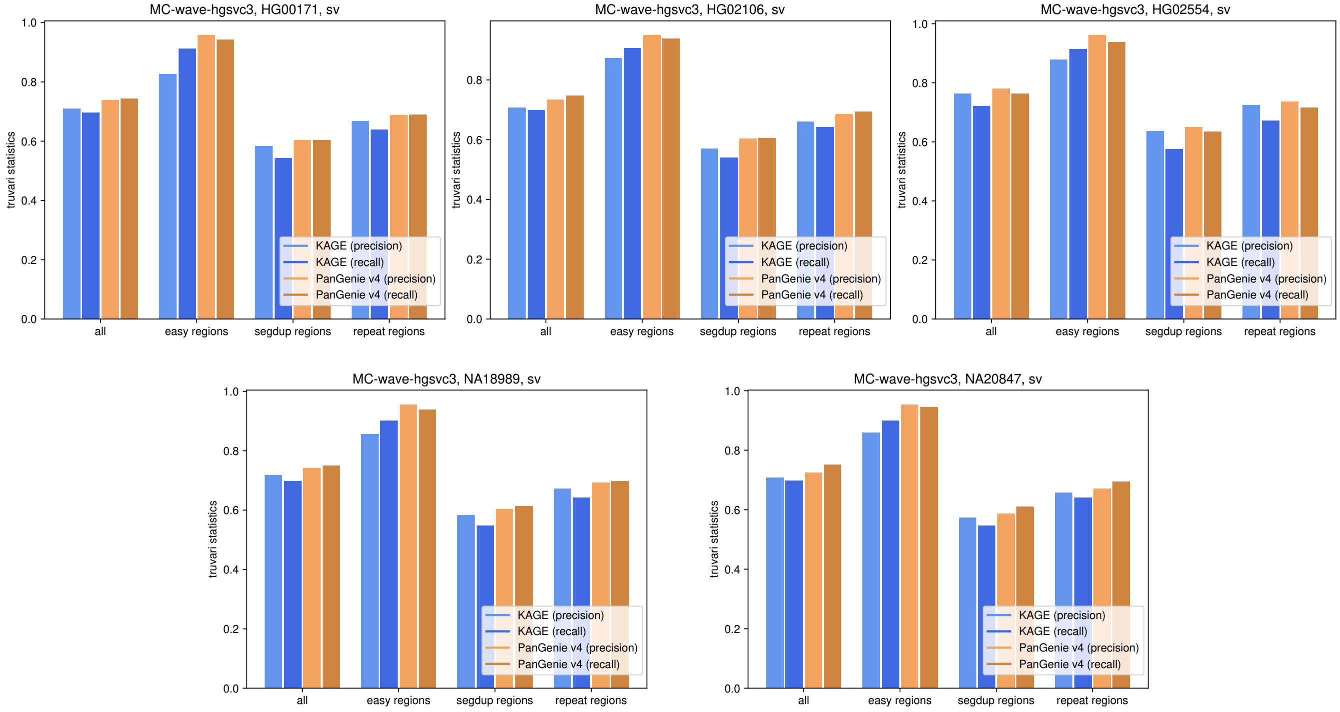


Figure 9: **Evaluation of SVs with MC ground truth** Precision and recall statistics for PanGenie and KAGE computed for five samples based on the HGSVC3 Minigraph-Cactus ground truth SV calls. Results are stratified by GIAB regions.

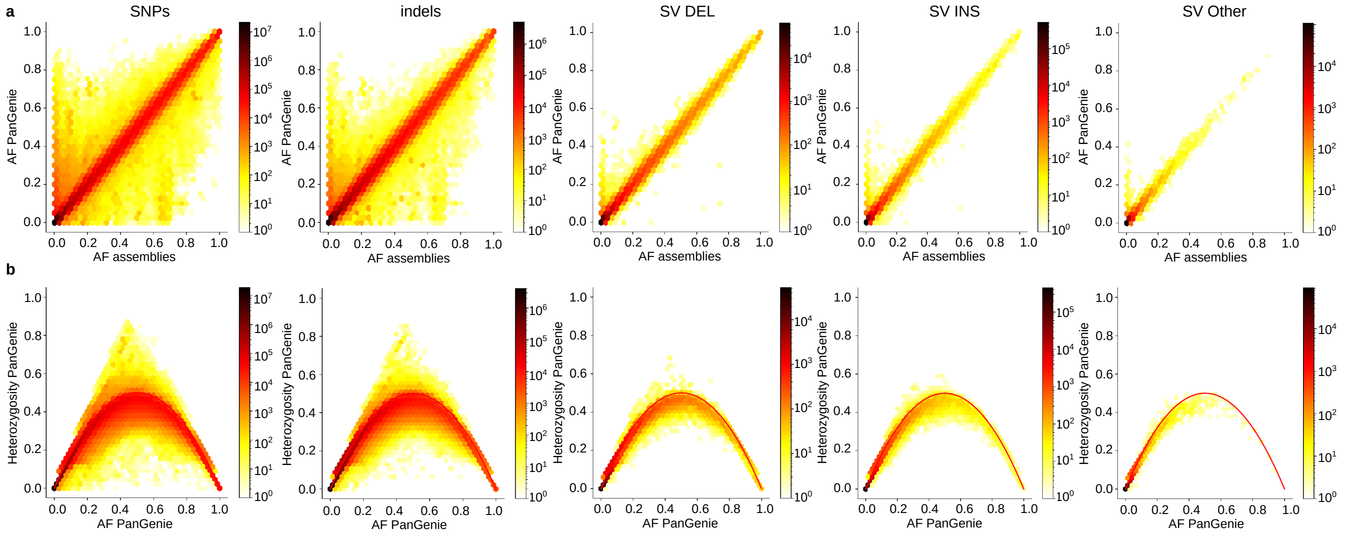


Figure 10: **Unfiltered PanGenie genotypes a** Comparison of allele frequencies of all SNPs, indels and SVs after genotyping across the 1kGP cohort with PanGenie to allele frequencies of the same variants in the assemblies. **b** Heterozygosities of SNPs, indels and SVs after genotyping across the 1kGP cohort with PanGenie as a function of the allele frequency.

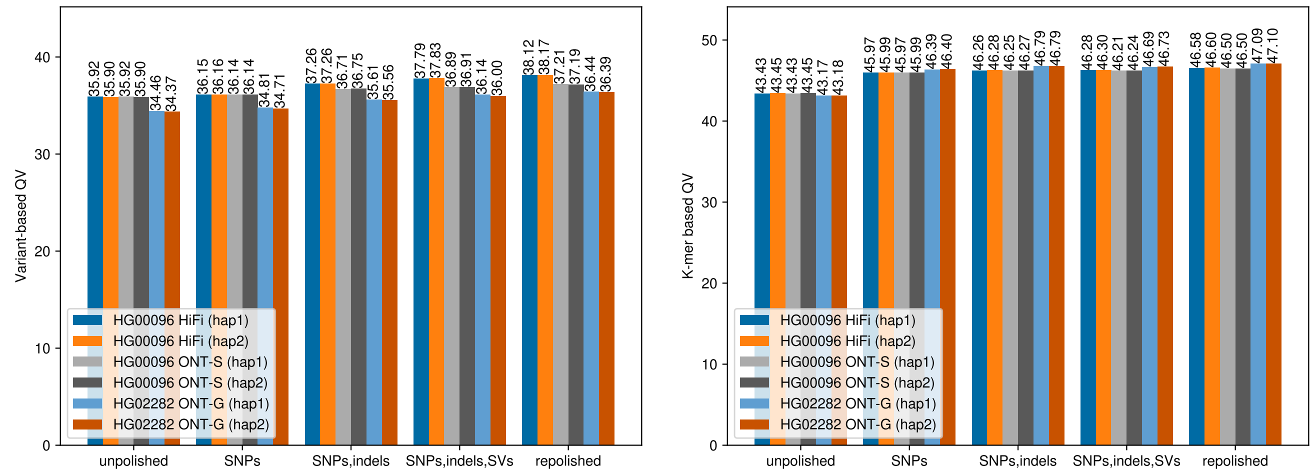


Figure 11: **Polishing performances** Variant-based QVs (left) and k-mer based QVs (right) computed for unpolished and polished haplotypes. Polishing was run on different long read datasets: ONT data produced by Schloissnig et al. (ONT-S) [12], ONT data produced by Gustafson et al. [13] and HiFi data [5]. Furthermore, different polishing runs were performed including only SNPs, SNPs and indels and all variant types (SNPs, indels, SVs). For the latter case, we also ran a repolishing experiment in which we polished haplotypes twice.

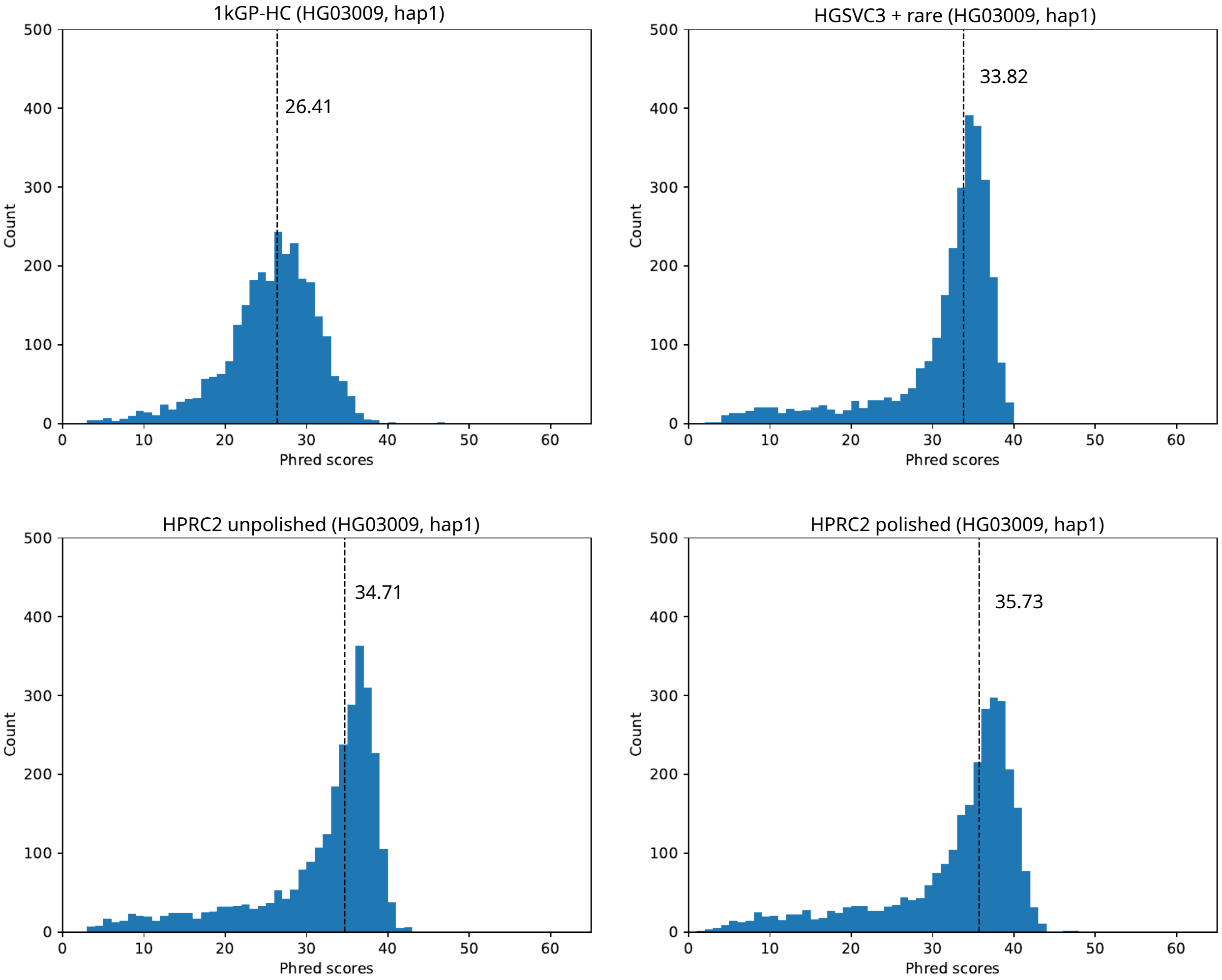


Figure 12: **Variant-based QV histograms for HG03009** Variant-based QV estimates were computed by comparing consensus haplotypes to ground truth HGSVC3 assemblies of the respective samples. QVs were computed based on the basepair changes observed across intervals of 1Mbp in length. The plots show the distribution of these QVs across all intervals. The median is marked by a dotted line. We evaluated consensus haplotypes computed based on four different datasets: the 1kGP phased variants called with traditional short-read based methods [6], the HGSVC3 PanGenie consensus haplotypes with added rare SNPs+indels [5], the unpolished HPRC v2.0 consensus haplotypes and the polished HPRC v2.0 consensus haplotypes. Note that, since HGSVC3 assemblies are used as a ground truth, sample HG03009 was contained in the input panel used to compute the HGSVC3 PanGenie genotypes.

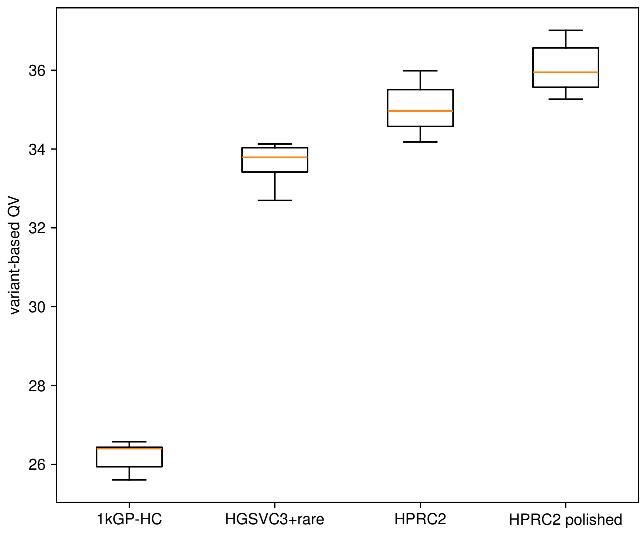


Figure 13: **Variant-based QVs** Variant-based QVs computed based on HGSVC3 assemblies across 8 evaluation samples. Variant-based QVs were computed for consensus haplotypes constructed from four different datasets: the 1kGP phased variants called with traditional short-read based methods [6], the HGSVC3 PanGenie consensus haplotypes with added rare SNPs+indels [5], the unpolished HPRC v2.0 consensus haplotypes and the polished HPRC v2.0 consensus haplotypes. For the 1kGP set, PAV failed on sample HG01596 and thus no variant-based QVs could be computed. Therefore, the corresponding boxplot contains results for 7 samples only.
